# Centrosomal Enrichment and Proteasomal Degradation of SYS-1/β-catenin Requires the Microtubule Motor Dynein

**DOI:** 10.1101/2021.11.10.468136

**Authors:** Joshua W. Thompson, Maria F. Valdes Michel, Bryan T. Phillips

**Affiliations:** University of Iowa

**Keywords:** Development, Wnt, β-catenin, microtubule, ECM29, dynein, proteasome, trafficking, elegans, asymmetry, cell cycle, robustness

## Abstract

The *C. elegans* Wnt/β-catenin Asymmetry (WβA) pathway utilizes asymmetric regulation of SYS- 1/β-catenin and POP-1/TCF coactivators. WβA differentially regulates gene expression during cell fate decisions, specifically by asymmetric localization of determinants in mother cells to produce daughters biased towards their appropriate cell fate. Despite the induction of asymmetry, β-catenin localizes symmetrically to mitotic centrosomes in both mammals and *C. elegans*. Due to the mitosis-specific localization of SYS-1 to centrosomes and enrichment of SYS-1 at kinetochore microtubules when SYS-1 centrosomal loading is disrupted, we investigated active trafficking in SYS-1 centrosomal localization. Here, we demonstrate that trafficking by microtubule motor dynein is required to maintain SYS-1 centrosomal enrichment, by dynein RNAi-mediated decreases in SYS-1 centrosomal enrichment and by temperature-sensitive allele of the dynein heavy chain. Conversely, we observe depletion of microtubules by nocodazole treatment or RNAi of dynein-proteasome adapter ECPS-1 exhibits increased centrosomal enrichment of SYS-1. Moreover, disruptions to SYS-1 or negative regulator microtubule trafficking are sufficient to significantly exacerbate SYS-1 dependent cell fate misspecifications. We propose a model whereby retrograde microtubule-mediated trafficking enables SYS-1 enrichment at centrosomes, enhancing its eventual proteasomal degradation. These studies support the link between centrosomal localization and enhancement of proteasomal degradation, particularly for proteins not generally considered ‘centrosomal’.

## Introduction

During metazoan development, cells must robustly respond to and integrate multiple signaling pathways to appropriately specify and populate tissues. This is of further importance in the developing embryo, where cell division rapidly alternates between S-phase and mitosis while simultaneously specifying tissues and entire germ layers (Pintard and Bowerman 2019). As such, the cell’s response to signals and the subsequent establishment of asymmetry often begins before daughter cells are ‘born’ via an asymmetric cell division (ACD) (Sunchu and Cabernard 2020; Juanes, 2020; Fuchs and Chen, 2013; Gómez-López, Petrisch et al 2014). Among the signaling pathways utilized in development is Wnt/β-catenin signaling, wherein the binding of Wnt ligands dismantlesthe Axin and APC containing destruction complex. This stabilizes transcriptional co-activator β-catenin, enabling it to activate target genes with activator TCF/LEF (Nusse and Clevers 2017). Wnt/β-catenin signaling is specialized in its ability to regulate both polarity and proliferation – with the Wnt signal orienting and promoting expansion of tissues simultaneously (Loh et al 2016).

Due to its role in both proliferation and polarity, Wnt/β-catenin can bewellused bythe organization to pattern a stem cell niche for tissue maintenance (Pinto, Clevers et al 2003; Reilein, Kalderon et al 2017). However, this function in adult tissue homeostasis also implicates dysregulation of Wnt/β-catenin signaling with hyper-proliferative disease states. Throughout transduction of the Wnt signal are proteins vulnerable to oncogenic mutation (Yang, yen 2011; Zhunussova, Djansugurova et al 2019; Liu, Thibodeau et al 2000). However, these often affect their tumorigenicity via dysregulation of the pathway effector β-catenin, which can promote proliferation by over-and mis-expression of targets like Cyclin D1 and cMyc or metastasis by disrupting adherens junctions (Niehrs, Acebron 2012; He, Kinzler et al 1998; Shtutman, Ben-Ze’ev et al 1999; Tetsu and McCormick, 1999; Reya, Clevers 2005).). The combination of these roles for β-catenin makes it an attractive target for studying tumorigenesis. However, the dual functions can also lead to confounding pleiotropic phenotypes after manipulation of β-catenin activity (Reya, Clevers 2005; Tejpar, Alman et al 1999; Geyer, Reis-Filho et al 2011).

Unlike β-catenin in the canonically described Wnt signaling pathway, *C. elegans* has distributed the adhesive and signaling roles across its four β-catenins (Sawa, 2012). Here we focus specifically on the *C. elegans* β-catenin SYS-1 for its role as a transcriptional co-activator in a modified version of canonical Wnt signaling, the Wnt/β-catenin Asymmetry pathway(WβA) (Phillips, Kimble et al 2007). Similar to canonical Wnt, the WβA pathway regulates co-activator SYS-1 (β-catenin), which modulates the activity of transcription factor POP-1 (TCF) during asymmetric cell division (Baldwin and Phillips 2018, Lam and Phillips 2017). Mother cells undergoing a WβA-driven division polarize components of the Wnt signaling pathway along the axis of the division – distributing positive regulators of a Wnt response, the Frizzled receptor and downstream effector Disheveled, to the signaled pole of the cell, and negative regulators, members of the β-Catenin destruction complex APC and Axin, to the unsignaled pole (Baldwin and Phillips 2014; Baldwin, Phillips et al 2016; Sugioka, Sawa et al 2011; Mizumoto and Sawa, 2007). Daughter cells in such a division therefore inherit Wnt signaling components that bias them toward their appropriate cell fate. Contrary to its known role asymmetrically activating Wnt target genes, SYS-1 (and indeed, mammalian β-catenin (Mbom, Barth et al 2014; Vora, Phillips et al 2020)), unexpectedly localizes symmetrically to mother cell mitotic centrosomes (Phillips, Kimble et al 2007; Huang, Lin et al 2007; Vora and Phillips 2015).

Previous work has demonstrated that centrosomal localization of SYS-1 in mother cells limits SYS-1 levels in daughter cells in a proteasome dependent manner (Vora and Phillips 2015). Similarly, differential localization of the proteasome has been shown to structurally and functionally disrupt protein complexes (Baldin, Coux et al 2008; Amsterdam, Baumeister et al 1993), suggesting that the regulation and localizationof protein clearance are interdependent. Proteasomal subunits have specifically been shown to co-sediment with centrosomal proteins and perinuclear inclusions, demonstratingthat the proteasome may be centrosomally localized (Wigley, Thomas et al 1999; Wojcik, DeMartino 2003). Similarly, turnover of the SYS-1 mobile fraction at mitotic centrosomes is enhanced by RPT-4 proteasomal activity (Vora and Phillips 2015). When the localization of SYS-1 to centrosomes is reduced by *rsa-2(RNAi)*, resultant daughter cells retain 40% additional SYS-1 protein. Perhaps, then, the centrosome is a region where SYS-1 and regulators of its degradation are concentrated to enhance SYS-1 regulation.

The directionality and temporal specificity of SYS-1 enrichment at only mitotic centrosomes implied that a more active localization mechanism might be at work, as compared to a passive diffusion-capture mechanism. Centrosomal SYS-1 maintains a proteasome-dependent, 100% mobile fraction, which is recycled within 120 seconds, yet remains undetectable via SYS-1 fluorescence on interphase centrosomes(Vora and Phillips 2015). The mechanism of such rapid and temporally specific SYS-1 localization is unknown. Several observations indicate that cells utilize the microtubule cytoskeleton for rapid and directional localization of cargo, including: transcription factors due to neuronal injury or signaling status, (Ben-Yaakov, Fainzilber et al 2012; Mikenberg, Kaltschmidt et al 2007) and endocytic recycling components around the Microtubule Organizing Center (MTOC) (Horgan, Mccaffrey et al 2010) or preventing pathogenic accumulation of RNA binding proteins (Deshiumaru, Tsuboi et al 2021). Therefore, we hypothesized that the role of the mitotic centrosome as an MTOC could facilitate the rapid transport of SYS-1 to the pericentriolar material specifically in dividing cells.

Accumulation of SYS-1 at an MTOC like the mitotic centrosome is most likely accomplished by the minus end-directed dynein microtubule motor complex (Vora and Phillips 2015; Priyanga, Bhaka-Guha et al 2021). The dynein motor is a multi-subunit complex composed of a pair of ATP-ase, microtubule-binding heavy chains responsible for the force-generating steps of the motor, and a variable complement of smaller light, intermediate, and accessory chains that alter cargo binding or the processivity of the motor (Reck-Peterson, Carter et al 2018; Jha and Surrey 2015). The dynein motor is multifunctional - it provides pulling forces, positions organelles, and can mediate rapid and conditional movement of proteins and RNAs (Carminati and Stearns 1997; Woźniak, Allan et al 2009; Harada, Hirokawa et al 1998; Hanz, Fainzilber et al 2003; Bullock and Ish-Horowicz 2001). For example, injured rat axons have demonstrated a dynein intermediate chain-dependent nuclear localization and coimmunoprecipitation of the transcription factor STAT3 to transcriptionally respond to injury (Ben-Yaakov, Fainzilber et al 2012). Dynein specifically is the motor complex most often directed towards the MTOC, and is known to be active during mitosis (Jha and Surrey 2015). Therefore, we focused our investigation on dynein-dependent trafficking as an ideal mechanism for SYS-1 localization to the MTOC mitotic centrosomes.

Here we demonstrate that microtubules and minus-end microtubule-mediated trafficking are required for both the centrosomal recruitment and clearance of SYS-1. We evaluate the extent to which SYS-1 utilizes dynein-mediated trafficking to accumulate at centrosomes by RNAi screen, and evaluate the effect of long and short term disruptions to microtubule-mediated trafficking on SYS-1 localization, target gene activity, and cell fate decisions.. Together, these data characterize a role for microtubule-mediated trafficking in arobust response to the WβA signaling pathway by enhancing the interaction of SYS-1 with negative regulators at the mitotic centrosome.

## Results

### SYS-1 Overexpression and Centrosomal Uncoupling Increases SYS-1 Localization to Regions of Dynein Enrichment

To test the role of microtubule-mediated trafficking in SYS-1 centrosomal localization, we investigated what trafficking mechanisms are necessary for SYS-1 regulation. As described above, we focused on the minus-end directed cytoplasmic dynein motor complex. The dynein localization pattern corresponded to that of GFP::SYS-1, localizing to centrosomal, cortical, and microtubule dense kinetochore regions (Schmidt, Van den Heuvel et al 2017; Tame, Medema et al 2014). Moreover, the kinetochore and cortical localization of GFP::SYS-1 was more common upon disruption of SYS-1 centrosomal localization by RNAi depletion of RSA-2(Fig. 1A).

**Figure 1:**
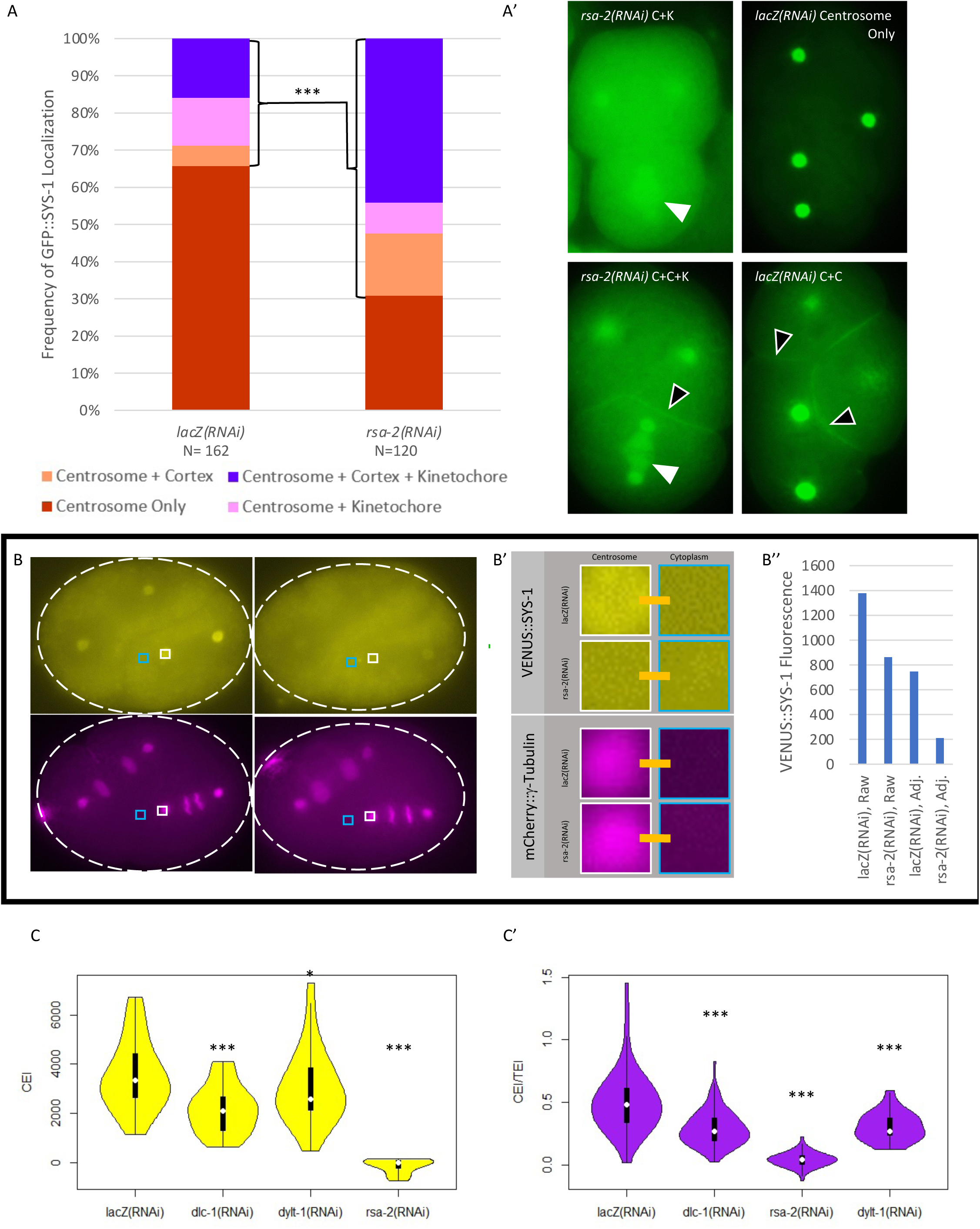
Rationale and methodology for microtubule motor RNAi Screen. A) GFP::SYS-1 can be occasionally observed at regions corresponding to DHC-1 mitotic accumulation. This is particularly enhanced in centrosomally uncoupled *rsa-2(RNAi)* embryos. The frequency of GFP::SYS-1 enrichment at these locales in lacZ*(RNAi)* and rsa-2*(RNAi)* treatment is distinguished by color. A’) GFP::SYS-1 at kinetochore microtubule (white arrowhead) and cortex(black arrowhead) of the dividing *C. elegans* early embryo. B) Demonstration of CEI and TEI calculation. CEI was calculated in FIJI by subtracting the mean pixel intensity of a cytoplasmic ROI encompassing the proximal half of the dividing cell, approximated by the light blue square, from a circular centrosome ROI, approximated by the white square. B’) Expanded view of ROI’s as defined in B. B’’) Graphical representation of the raw centrosomal and CEI adjusted fluorescence intensity for the control (Left) and Affected (right) embryos in B. Control (*lacZ(RNAi)* treated) exhibits both a high raw centrosomal GFP::SYS-1 and CEI. Affected (*rsa-2(RNAi)* treated) exhibits an intermediate raw GFP::SYS-1, but severely decreased CEI, better reflecting its visual failure to increase centrosomal GFP::SYS-1 notably above the bulk of cytoplasmic representation. C) Example VENUS::SYS-1 CEI (C) and CEI/TEI (C’) of controls and indicated dynein light chain depletions. Width of the violin plot corresponds to a histogram of observation density. The median is indicated by the central white dot, with the interquartile range indicated by the thick black line, and 1.5x additional Interquartile range indicated by the thin black line. N= lacZ*(RNAi)*, 72; *dlc-1(RNAi),* 74; *dylt-1(RNAi),* 52; *rsa-2(RNAi)*, 22 * - p<0.05, *** - p<0.001 by Chi-square test (A) or Student’s t-test (C-C’).

While GFP::SYS-1 has been observed at mitotic centrosomes in many dividing cells throughout development, localization to these cortical and kinetochore ‘secondary’ sites is uncommon (accumulation at either or both of these sites occurring in 34.5% of embryos) in untreated embryos (Fig.1A). Uncoupling SYS-1 protein from the centrosome by *rsa-2(RNAi)*, leads to more frequent detection of SYS-1 at sites of dynein and kinesin localization (Fig. 1A). Given its comparable localization to kinetochore microtubules, cortex, and the MTOC/centrosome, SYS-1 may be a cargo of the dynein or kinesin microtubule motors (Vora and Phillips 2015; Vora and Phillips 2017; Ferenz, Wadsworth et al 2010; Schmidt, Van den Heuvel et al 2017).

### Knockdown of Dynein Complex Subunits Reduces SYS-1 Centrosomal Localization

Given the observation that SYS-1 localizes to cortical and kinetochore sites that also exhibit enrichment for microtubule motors, we investigated a functional role for microtubule-mediated trafficking in SYS-1 localization. We began by performing an RNAi screen against components of the dynein motor complex and dynein associated proteins, expanding to additional motors and known microtubule-affecting proteins (Fig. S1). We assayed the centrosomal enrichment of GFP::SYS-1 in each of these depletions. We expected depletion of transport proteins that play a role in SYS-1 centrosomal transport to be defective in the maintenance of SYS-1 centrosomal enrichment when SYS-1 centrosomal clearance was otherwise unaffected. In order to evaluate the centrosome-specific enrichment of SYS-1, we considered two ways of comparing the extent of fluorescently-tagged SYS-1 localization to centrosomes: 1) the centrosomal enrichment index (CEI,similar to that described in Gao, Smith et al 2017), the centrosomal fluorescence intensity with background cytoplasmic fluorescent intensity of the surrounding cell subtracted, to control for backgrounds that may pleiotropically influence SYS-1 expression, and 2) the SYS-1 CEI evaluated proportionally to the enrichment factor of representative Peri Centriolar Material (PCM) factor γ-tubulin (Tubulin Enrichment Index, or TEI), to avoid false positives via defects in PCM maturation (CEI/TEI) (Fig. 1B-B’’). We then compared these centrosomal enrichment metrics between microtubule motor depletions and established *rsa-2(RNAi)* centrosomal uncoupling (Fig. 1C-C’).

As expected, depletion of dynein subunits often resulted in pleiotropic and deleterious effects, including oogenesis and early patterning defects that precluded our SYS-1 localization assay. We therefore treated animals with partial RNAi knockdowns by commencing RNAi feeding 12-24 hours before imaging as necessary, based on the phenotypic severity. While no treatment disrupted SYS-1 centrosomal levels as severely as *rsa-2(RNAi)*, we did observe several statistically significant decreases in centrosomal SYS-1 enrichment (Fig. 21C – C’, Fig. S1). Furthermore, while the total population of some treatments displayed only an intermediate decrease in SYS-1 CEI, the distribution of SYS-1 CEI values observed in dynein subunit knockdowns extends from wild-type to an *rsa-2(RNAi)-*like disruption (e.g., Fig. 2 Supplement 1). Dynein knockdown-induced decreases in SYS-1 CEI or CEI/TEI were therefore masked somewhat by their broad range in phenotype and/or RNAi penetrance (Fig. 1C-C’, Fig S2). Most notably, *dlc-1(RNAi)* and *dylt-1(RNAi)* each exhibit a significant reduction in centrosomal SYS-1 localization disproportionately larger than the effect on γ-Tubulin. Loss of centrosomal SYS-1 enrichment was specifically more visible in a transgene driven by the endogenous SYS-1 promoter (P_sys-1_::VENUS::SYS-1), as compared to SYS-1 driven by the stronger *pie-1* promoter (P_pie-1_::GFP::SYS-1) (Fid 2C-C’).

**Figure 2:**
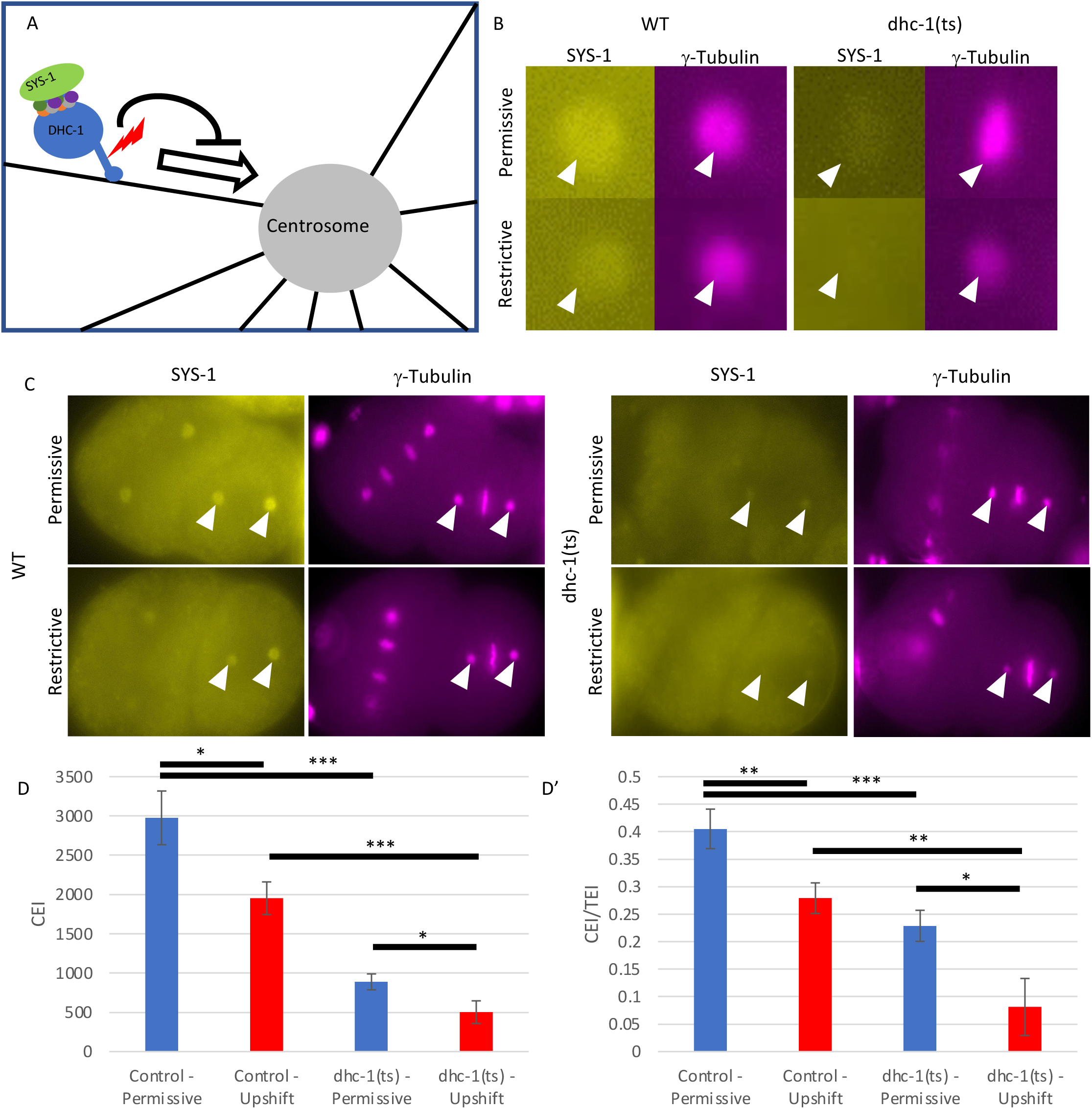
The dynein heavy chain Temperature Sensitive (dhc-1(ts)) Allele mediated loss of dynein function has a disproportionate effect on SYS-1 localization. A) Diagram of the Dynein motor, and hypothetical model of its role in SYS-1 trafficking. Relative location of *dhc-1(ts)* lesion indicated. B) Representative centrosome images of VENUS::SYS-1 and g-Tubulin::mCherry in wildtype and ts allele-containing embryos C) embryos at comparable cell cycle phase before (top) and after (bottom) temperature upshift. D) SYS-1 CEI and D’) SYS-1 CEI per TEI in embryos containing *dhc-1(ts)*, to control for differences in centrosome assembly, standard error depicted. N= 44, 31, 38, 25 in D and D’ columns, left to right. * - p<0.05, ** - p<0.01, *** - p<0.001 by Student’s t-test.

While we observed dynein-dependent decreases in SYS-1 localization, this could be because of changes in the RSA-2 centrosomal scaffold, which is responsible for capturing SYS-1 at mitotic centrosomes (Vora and Phillips 2015, Schlaitz, Hyman et al 2007). Depleting some dynein subunits did result in a slight decrease in centrosomal GFP::RSA-2 - 16% reduction in dlc-1(RNAi) embryos and 14% in dylt-1(RNAi) embryos. Embryos depleted of DLC-1 by RNAi exhibit a much more pronounced effect on SYS-1 localization, with 39% and 42% reductions in P_sys-1_::VENUS::SYS-1 CEI and CEI/TEI, respectively. DYLT-1 depletion exhibited a similar 14% decrease in CEI alone, but a 39% decrease in CEI/TEI. Both results suggest that tagged SYS-1 is disproportionately removed from centrosomes in these dynein subunit knockdowns (Fig. S3)

These data suggest that dynein does have a role in the localization of SYS-1 to centrosomes, but redundancy between components of the dynein motor or between the dynein motor and other trafficking processes may limit the effect any one subunit has on SYS-1 localization. Alternately, variability of RNAi efficacy may have resulted in incomplete depletion of SYS-1 relevant dynein subunits. Similarly, the pleiotropic effects of dynein depletion may also have precluded our ability to measure their effects on SYS-1 localization, leading to the evaluation of only partial knockdowns. In order to address these issues, we investigated the role of the dynein motor in SYS-1 localization at a more precise temporal resolution via conditional dynein disruption.

### Temporally Restricted Dynein Loss of Function Reveals Dynein Requirement for Centrosomal SYS-1 Accumulation

While the results of our screen demonstrate that dynein knockdown significantly reduces SYS-1 centrosomal enrichment, rare individuals exhibiting *rsa-2(RNAi)-*like loss of centrosomal SYS-1 accumulation, we were unable to completely recreate a population exhibiting consistently severe disruption to SYS-1 CEI. To distinguish between functional redundancies limiting the effects of dynein depletion and the confounding deleterious effects described above, we utilized a temperature sensitive allele of the cytoplasmic dynein heavy chain, *dhc-1(or195ts),* to conditionally inactivate the cytoplasmic dynein heavy chain. This allele, identified in the Bowerman lab at the University of Oregon, introduces a lesion to the microtubule binding stalk of DHC-1. This lesion renders the motor slightly hypomorphic at the permissive 15°C (with 91% of embryos remaining viable and fertile) but complete mitotic spindle collapse occurs in as little as 30 seconds upon upshift, suggesting that DHC-1 motor function has been nearly completely ablated (O’rourke, Bowerman et al. 2007).

We expected that combining the *dhc-1(ts)* allele with fluorescently tagged SYS-1 would allow us to examine a more penetrant dynein knockdown. This allowed us to overcome both redundancy between dynein accessory chains in SYS-1 trafficking as well as the previously described incomplete or pleiotropic effects of dynein depletion (Fig. 2A). We compared the CEI of P_sys-1_:VENUS::SYS-1 (hereafter VENUS::SYS-1) to that of mCherry::γ-Tubulin (TEI). SYS-1 and γ-Tubulin are both recruited to centrosomes at a much higher rate during mitosis than in interphase (Vora and Phillips 2015; Raynaud-Messina and Merdes 2007). Because *dhc-1(ts)* limits the extent of γ-Tubulin recruitment (Fig. 2B-C), evaluating SYS-1 CEI as a ratio to g-Tubulin TEI ensures that decreases to the SYS-1 CEI must be disproportionately large to be numerically distinct.

Consistent with the published hypomorphic nature of the allele at the permissive temperature (O’rourke, Bowerman et al 2007), *dhc-1(ts)* embryosexhibit a notable decrease in the SYS-1 CEI of embryos kept at the permissive temperature (Fig. 2 B-D). Following 30-45 seconds of temperature upshift, the hypomorphic SYS-1 localization is enhanced, with severely reduced SYS-1 centrosomal enrichment (Fig. 2B-C). While there was a decrease in centrosome size, presumably due to roles for dynein in centrosome maturation (Priyanga, Bhakta-Guha et al 2021), we accounted for this by assessing proportional changes in SYS-1 CEI compared to changes in γ-Tubulin TEI. We observed a CEI/TEI decrease, corresponding to decreased centrosomal SYS-1 accumulation, proportionally greater than the reduction in γ-tubulin (Fig. 2D’). When compared to the individual dynein subunit knockdown, the more penetrant phenotype of *dhc-1(ts)* mutants indicates that SYS-1 requires the dynein microtubule motor complex to localize appropriately to centrosomes. Additionally, it seems likely that redundancy of peripheral motor components or incomplete knockdown is indeed responsible for the variability within individual subunit depletions seen above (Fig 1C-C’).

### Microtubule Depolymerization Experiments Reveal that SYS-1 Centrosomal Dynamics are Affected both Positively and Negatively by Microtubule-Mediated Trafficking

To determine the extent to which microtubule-based trafficking in general is responsible for SYS-1 centrosomal localization, we treated gravid adults and their embryos with >100 μM nocodazole, a microtubule-destabilizing agent. This dose, while sufficient for a rapid acute response in adult animals, resulted in either no effect on fertilized embryos because of the eggshell or prevented the production of additional embryos via germline arrest (unpublished data). We therefore turned to a lower dose, longer-term nocodazole treatment: 12 hours at 25μg/mL (83 μM). Maternal nocodazole treatment allowed us to obtain embryos with limited microtubule networks, but sufficient spindle development to enter mitosis (Fig. 3A). Surprisingly, the centrosomal enrichment of SYS-1::GFP driven by a general maternal promoter (P_pie-1_::GFP::SYS-1, hereafter GFP::SYS-1) increased after nocodazole treatment (Fig. 3B, C).

**Figure 3:**
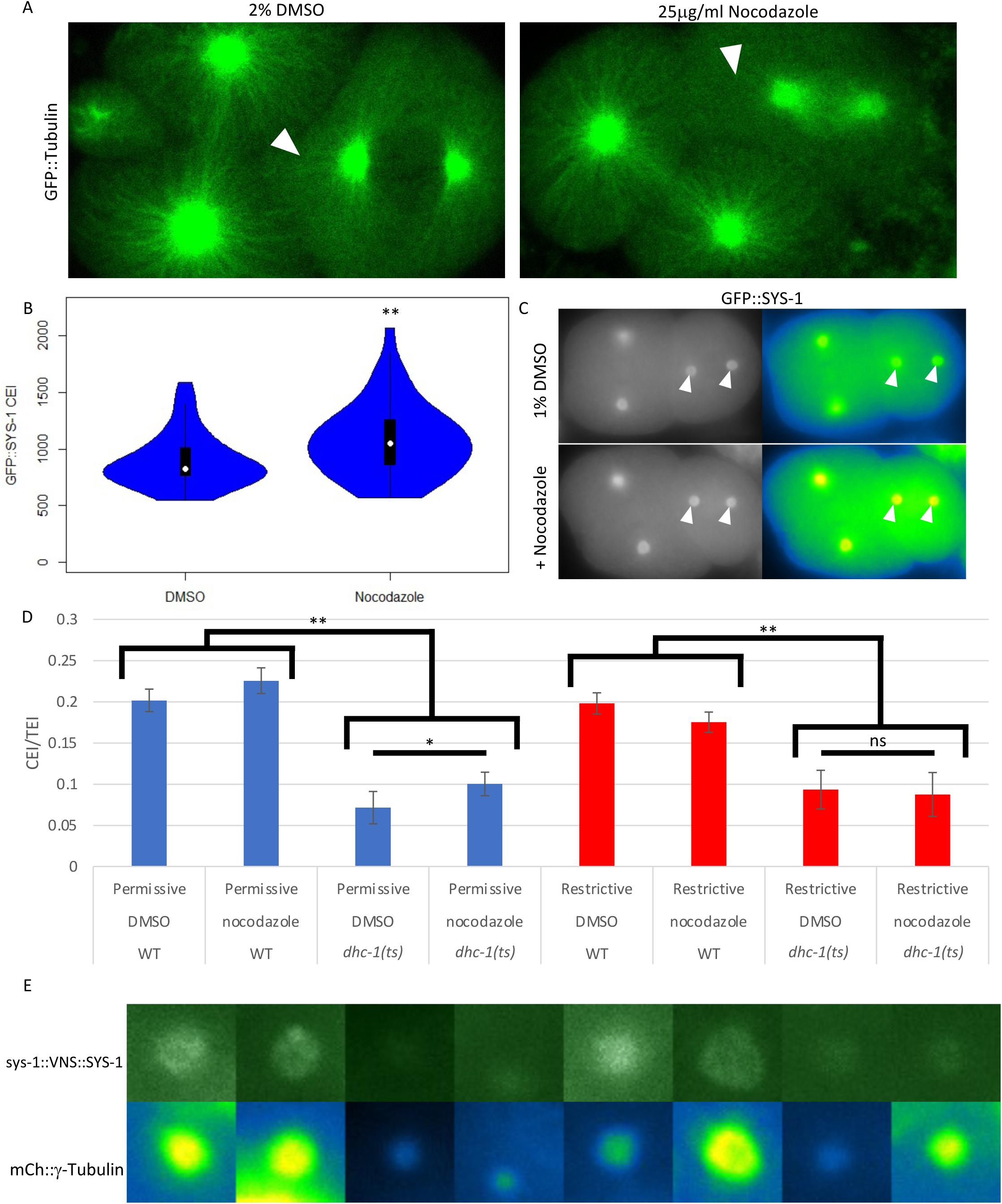
Nocodazole mediated disruption of microtubules enhances centrosomal GFP::SYS-1 enrichment. A) Representative images of GFP::TBB-1 in DMSO and nocodazole treated embryos. White arrowheads indicate astral microtubules (left) and the corresponding lack or dysregulated region in nocodazole treatment (right) B) Quantification of, and C) Representative images of similarly treated GFP::SYS-1 embryos. For B, N=36 DMSO and 46 nocodazole treated embryos. * - p<0.05, ** - p<0.01 by Student’s t-test. D) CEI/TEI as evaluated with combinatorial Wildtype (WT) or *dhc-1(ts)* allele for Dynein Heavy Chain, and treatment with 2% DMSO vehicle media or 2% DMSO + nocodazole media. Measurements at the permissive and restrictive temperature are differentiated by blue and red coloring, respectively. Error bars are SEM. E) Representative P_sys-1_VENUS::SYS-1 and mCh::g-Tubulin for each treatment in D, respectively.

The observed SYS-1 CEI increase in nocodazole treated embryos is similar to the phenotype observed in RPT-4 depletion, which decreases proteasome function (Vora and Phillips, 2015). To ensure that this change in SYS-1 localization was not because of off-target effects, we compared maternal feeding to that of *perm-1(RNAi)* permeabilized embryos. When treated with 10.5 μM nocodazole, these embryos formed sufficient microtubule networks to enter mitosis and form spindles, but these spindles were often inappropriately rotated or positioned (61% of imaged embryos, N=11/18), suggesting that astral microtubules have been severely disrupted (Fig. S4A). We observed a similar CEI increase in GFP::SYS-1 in these embryos (Fig. S4B). This could be consistent with a model where both SYS-1 recruitment to centrosomes and SYS-1 proteasomal degradation are affected by disruptions to microtubule mediated trafficking. In this model subtle disruptions to dynein trafficking would limit the ability of SYS-1 to replenish its mobile fraction, reducing SYS-1 centrosomal enrichment (Phillips and Vora 2015). However, more severe disruptions to dynein function could be sufficient for defective localization of the regulatory process, preventing clearance of SYS-1 and increasing its centrosomal enrichment.

As different subsets of dynein and microtubule depletions exhibit either increased or decreased SYS-1 CEI, we investigated a combination of these treatments to observe the behavior of SYS-1 upon further disruption to microtubule-mediated trafficking. While both temperature upshift for *dhc-1(ts)* and nocodazole treatments significantly affect SYS-1 centrosomal localization, neither treatment appears to completely disrupt microtubule mediated trafficking. Nocodazole-treated embryos still exhibit some kinetochore microtubules (Fig. 3A) but can tolerate weakly poisoned microtubules for some time. Conversely, *dhc-1(ts)* embryos demonstrate severe loss of dynein function when reared at the restrictive temperature. Thus short-term temperature upshifts are required to establish spindles and maintain viability, suggesting that some portion of centrosomal trafficking during mitosis still occurs.”. We therefore tested the combination of these treatments.

In contrast to embryos with GFP:SYS-1 driven by the strong early embryonic PIE-1 promoter (Fig. 3 B-C), embryos containing P_sys-1_::VENUS::SYS-1, driven by the weaker endogenous SYS-1 promoter, exhibit only a slight, non-significant, increase in CEI/TEI when treated with nocodazole (Fig. 3D, 1^st^ and 2^nd^ columns). However, when P_sys-1_::VENUS::SYS-1 centrosomal enrichment is challenged by the hypomorphic *dhc-1(ts)* at the permissive temperature, nocodazole treatment is sufficient to significantly increase SYS-1 centrosomal recruitment. The finding that sensitizing embryos for dynein trafficking defects is required to observe a nocodazole phenotype for P_sys-1_::VENUS::SYS-1, but not for P_pie-1_::GFP::SYS-1 indicates that the difference in transgene expression uncover a differential nocodazole sensitivity in centrosomal recruitment of SYS-1. However, nocodazole-mediated increase in P_pie-1_::GFP::SYS-1 centrosomal enrichment is no longer observed when the nocodazole treated *dhc-1(ts)* embryos are upshifted to the restrictive temperature, suggesting that the increased SYS-1 centrosomal enrichment seen in nocodazole treated embryos still requires dynein trafficking.

These observations describe a centrosomal relationship wherein treatments limiting the coverage of microtubule-mediated dynein trafficking (i.e. nocodazole treatment) increase centrosomal SYS-1 because of relatively inefficient or rare trafficking events for SYS-1 negative regulators, while transient severe disruptions to dynein trafficking processivity (dynein subunit RNAi depletion, *dhc-1(ts)* temperature upshift, upshift + nocodazole) reveal that SYS-1 centrosomal enrichment requires continual replenishment via a functional microtubule motor.

If the nocodazole induced SYS-1 CEI increase described above is indeed due to a role for microtubule-mediated trafficking in the centrosomal enrichment of some component of the SYS-1 degradation pathway, we would expect this proteolytic activity to be similarly mislocalized upon trafficking disruption. Therefore, we decided to test the idea that proteasomal trafficking to, and enrichment at, the centrosome was also perturbed in nocodazole-affected microtubule networks. We first attempted to localize the proteasome. While the centrosome has been demonstrated to exhibit proteolytic activity (Vora and Phillips 2015; Vora and Phillips 2016; Hames, Fry et al 2005; Peel, O’Connell et al 2012; Wigley, Thomas et al 1999; Fabunmi, DeMartino et al 2000), existing immunohistochemical reagents for mammalian proteasome subunits did not display any distinct localization pattern in our embryos. An mCherry tag on proteasome subunit RPT-1 did demonstrate a kinetochore microtubule enrichment, similar to that of DHC-1 and SYS-1, but appeared absent from centrosomes. Inability to localize tagged RPT-1 may be due to a relatively low RPT-1 copy number per proteasome, or correspondingly low proteasome number, limiting its relative enrichment. Alternatively, fluorescently tagged proteins have shown evidence of degradation of the tag at a site of active proteolysis, particularly if such a tag limits proteasome activity (Huang, Lobel et al 2014). To circumvent this issue, we assayed for centrosomal proteasome activity indirectly by functionally testing an additional centrosomal target of the RPT-4 containing proteasome, the centriole duplicating kinase ZYG-1 (O’Connell, White et al 2001; Peel, O’Connell et al 2012). We additionally assayed therole of ECPS-1, the *C. elegans* homolog of mammalian ECM29, which is an adaptor protein that enhances local proteolytic activity of proteasomes via its function linking the dynein motor complex and the proteasome and regulating 26S proteasome distribution (Gorbea, Rechsteiner et al 2004; Gorbea, Rechsteiner et al 2010; Hsu, Cheng et al 2015; Ibañez-Vega, Yuseff et al 2021).

Knockdown of ECM29 homolog ECPS-1 (formerly D2045.2) resulted in embryos that displayed an average 33.4% increase in centrosomal P_pie-1_::GFP::SYS-1 enrichment, similar to both nocodazole-treated and *rpt-4(RNAi)* embryos (Fig. 4A, Fig. 3A-C). Therefore, the overall increase in SYS-1 CEI seen in nocodazole and *ecps-1(RNAi)* treated embryos suggests that microtubule mediated trafficking enhances the removal of SYS-1 from centrosomes as well. Given the role of ECM29 in regulating the distribution of proteasome complexes via microtubule-mediated trafficking, these data suggest that an improperly localized proteasome in ecps-1(RNAi) results in dysregulation of SYS-1 similar to that seen in proteasome functional knockdown (rpt-4(RNAi)) or broader depletion of microtubules (nocodazole treatment). To determine if the effects of ECPS-1 knockdown are specific to SYS-1 dysregulation, or may be due to more widespread effects on centrosomally localized proteins, we assayed an additional target. The ZYG-1 protein, a regulatory kinase controlling centriolar duplication, is both centrosomally enriched during mitosis and negatively regulated by the RPT-4 proteasome (Peel, O’Connell et al 2012). Utilizing a fluorescently tagged ZYG-1 fragment demonstrated to mirror the localization and dynamics of the endogenous protein (Peters, O’Connell et al 2010), we demonstrated 20% and 27% increased accumulation of ZYG-1 in both proteasome knockdown *rpt-4(RNAi)* and *ecps-1(RNAi)* treatments, respectively. The similarity of phenotype between nocodazole, ECPS-1 depletion, and RPT-4 proteasome knockdown implies a functional link between microtubule-mediated trafficking of centrosomally localized proteins and their eventual degradation by the RPT-4 proteasome.

**Figure 4:**
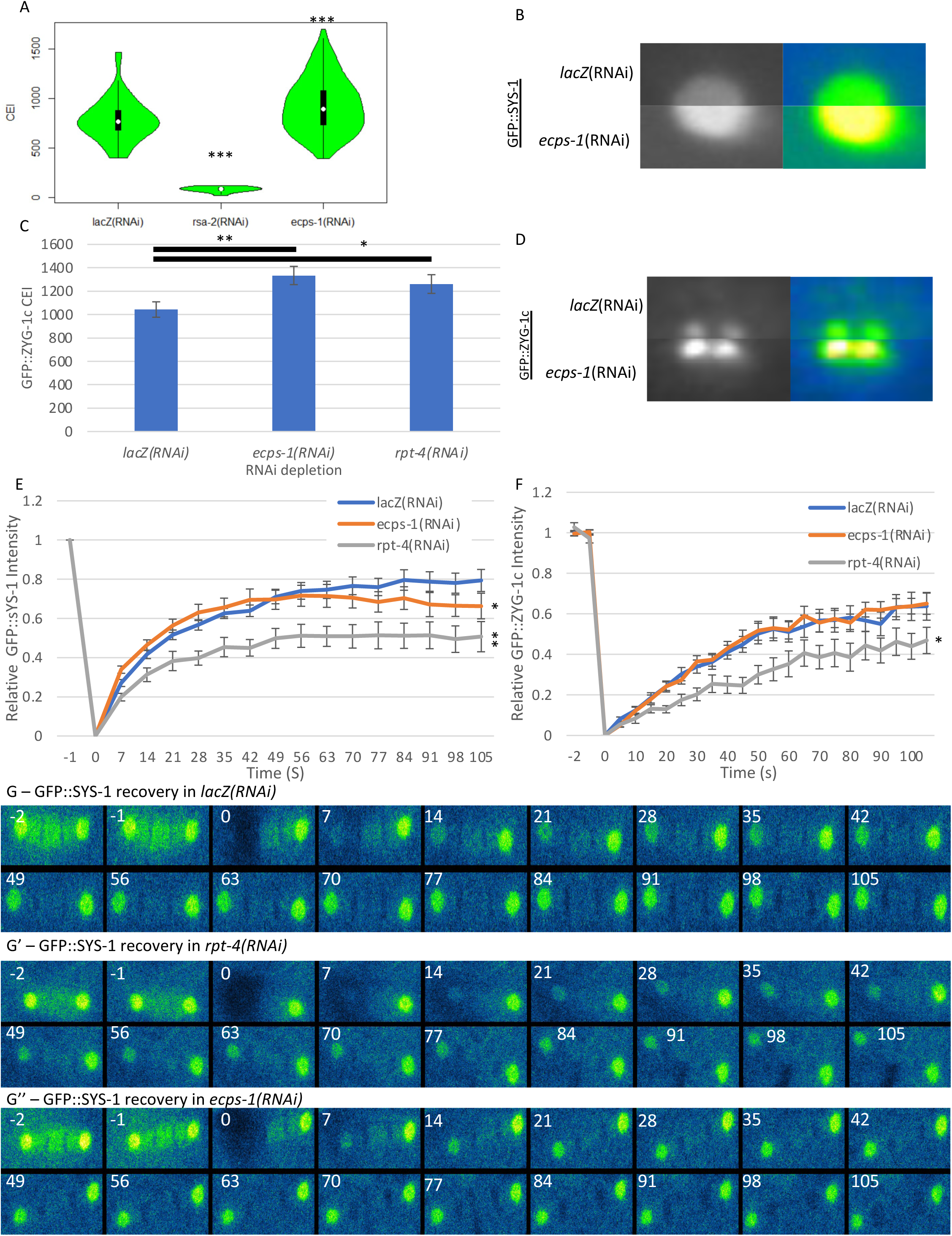
Microtubule-mediated processes can also stabilize centrosomal SYS-1. A) Treatment with *ecps-1(RNAi)* enhances centrosomal SYS-1 accumulation, N=48 *lacZ(RNAi), N =*12 *rsa-2(RNAi), N=* 124 *ecps-1(RNAi).* B) *lacZ(RNAi)* and *ecps-1(RNAi)* representative centrosomes in grayscale and intensity dependent look up table (LUT). C) Putative dynein and proteasome adaptor ECPS-1 also appears to also enhance centrosomal ZYG-1 accumulation, by GFP::ZYG-1c CEI (N=31 untreated, N=19 *ecps-1(RNAi),* N= 20 *rpt-4(RNAi),* 14 *rsa-2(RNAi)*) and D) Representative GFP::ZYG-1c centrosomal images in grayscale and intensity dependent LUT. (E-F) Quantified recovery of GFP::SYS-1 (E) and GFP::ZYG-1c (F). Values indicated are the ratio of bleached centrosomes to their prebleach value, normalized such that the postbleach value, t = 0, is 0 GFP::SYS-1 intensity. Error bars are SEM. (G–G’’) Representative FRAP montage of 2nd division P1 cells in the indicated embryos. Two prebleach, immediately consecutive time points are followed by the 1st postbleach image at time 0 and images every 7 s thereafter indicated at the top left. *p < 0.05, **p < 0.01, ***p < 0.001 by Student’s t test. One-sided Student’s t test, asterisk indicates p value from lacZ(RNAi) in E and F.

Because SYS-1 centrosomal recruitment depends on rapid degradation via the centrosomal proteasome (Vora and Phillips 2015), we assayed the mobile fraction of centrosomal SYS-1 in *ecps-1(RNAi)* by FRAP. Comparison of GFP::SYS-1 recovery after photobleach demonstrated that depletion of ECPS exhibits an intermediate phenotype between *rpt-4(RNAi)* proteasome depletion and negative controls (Figure G-G”). While the initial rate of *ecps-1(RNAi)* GFP::SYS-1 recovery was similar to that seen in wild type embryos, the late recovery phase of *ecps-1(RNAi)* embryos showed an early SYS-1 recovery plateau, evidence of a significantly reduced mobile fraction that was intermediate to wild type and proteasome depleted *rpt-4(RNAi)* embryos (Fig. 4E,G). The recovery curve observed in *ecps-1(RNAi)* suggests that SYS-1 recovers at an approximately wild type rate in *ecps-1(RNAi)* embryos but remains approximately as immobile as in proteasome-depleted embryos. Therefore, ECPS-1 depleted embryos appear to degrade SYS-1 incompletely, as observed in the RPT-4 proteasome depletion, but with sufficient protein turnover to enable wild type recruitment of newly trafficked SYS-1.

While both P_pie-1_:SYS-1::GFP and P_pie-1_::GFP::ZYG-1c exhibit an increase in their CEI as a result of *ecps-1(RNAi)*, their turnover rates at the centrosome are not similarly affected. This may be due to diverging regulatory pathways en route to the RPT-4 proteasome, or differences in the availability of binding sites at the centrosome specific to each protein. Meanwhile, ECPS-1 depleted embryos may be unaffected in protein clearance per proteasome, but the delayed or partial localization of the centrosomal proteasome could result in hierarchical stabilization of a spatially or temporally inaccessible fraction consisting of highly expressed and continually trafficked proteins, like SYS-1.

### Dynein Trafficking is Required for Restriction of Wnt-Signaled Cell Fates

Having established a two-fold role for SYS-1 regulation in microtubule-mediated trafficking, we investigated the role microtubule trafficking played in regulating SYS-1 dependent cell fate decisions. Since the SYS-1 centrosomal localization pattern has been seen across all examined SYS-1-expressing tissues (Phillips, Kimble et al 2007; Huang, Lin et al 2007; Baldwin and Phillips 2018; Lam and Phillips 2017), we investigated cell fate changes in embryonic and larval tissues across one embryonic and two larval SYS-1 dependent cell fate decisions. These included a transcriptional reporter of the *end-1* WβA target gene in the establishment of SYS-1 dependent endoderm, and anatomical markers for larval specification of the stem cell-like hypodermal seam cells and the germline stem cell niche.

First, to directly assay the effect of centrosomal dysregulation of transcriptional activator SYS-1, we investigated the nuclear enrichment of GFP::SYS-1 after division of the Wnt-regulated endomesodermal precursor cell (EMS)(Bei, Mello et al 2002). In the EMS division, SYS-1 is enriched in the posterior E nucleus, but not that of the anterior MS (Fig. 5A-B). Analyzing nuclear enrichment of green fluorescence protein (GFP)-tagged SYS-1, we were able to observe increased nuclear GFP signal in both the E and MS daughters in analyzed trafficking mutants (Fig. 5C). Interestingly, daughters affected by both *ecps-1(RNAi)* and *dlc-1(RNAi)* exhibit significant enhancement of nuclear enrichment in each of the two daughter cells (Fig 5C), suggesting that these treatments act negatively on SYS-1, despite opposing effect on SYS-1 centrosomal enrichment. Following loss of negative centrosomal regulation, these data suggest that daughter cell SYS-1 inheritance is increasing and thus affecting the bias of the EMS asymmetric division, increasing the likelihood of unsignaled daughter cells adopting the signaled endoderm cell fate.

**Figure 5:**
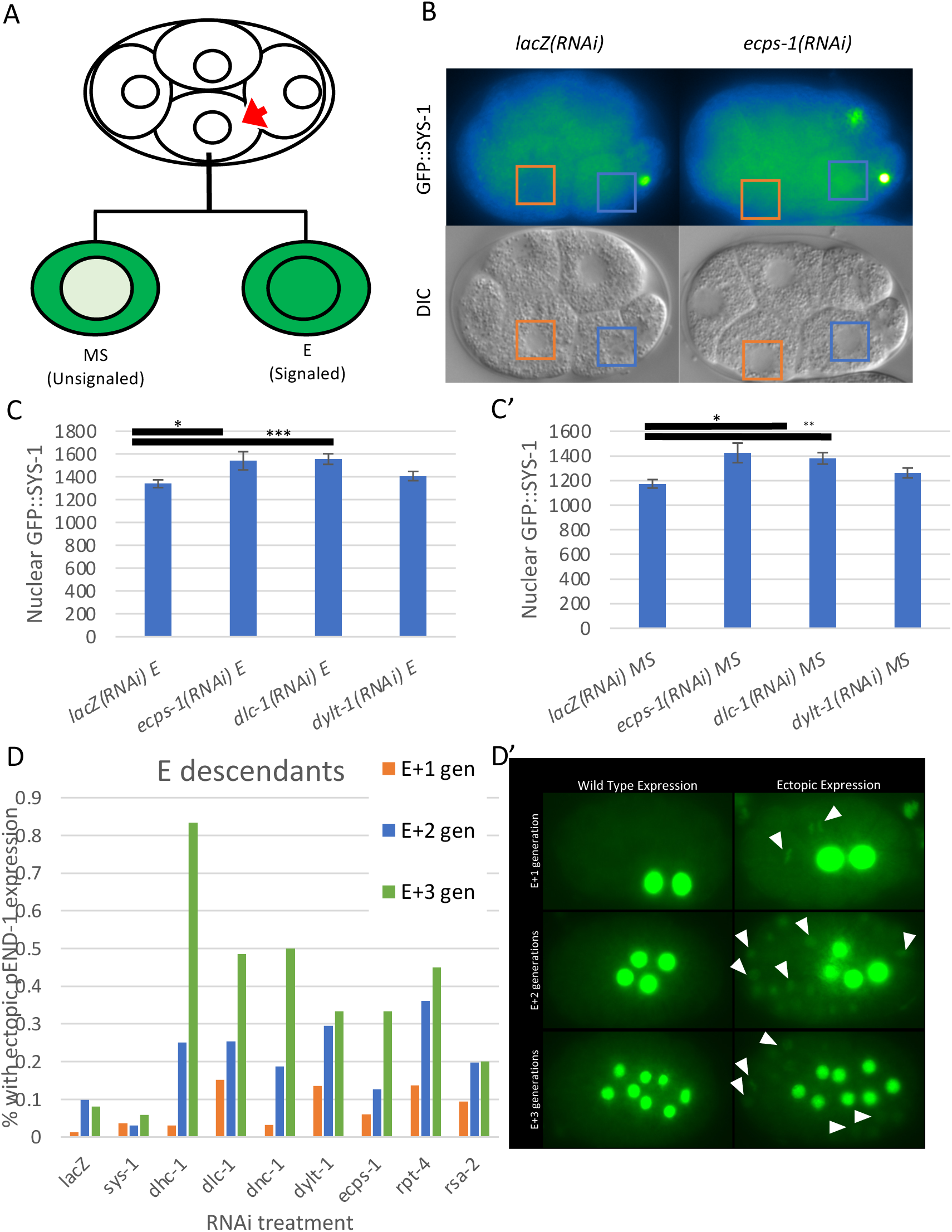
Effect of Dynein Depletion on nuclear SYS-1 Localization and Gene Expression in EMS Descendant Cells. A) The EMS precursor (embryo ventral cell) divides, receiving Wnt and Src specification and orientation cues, respectively (red arrow). These daughters give rise to posterior, signaled E and anterior, unsignaled MS daughter cells. Wildtype specification results in depicted approximate GFP::SYS-1 localization pattern specifically enriched in the signaled daughter nucleus. B) Representative images of wildtype and *ecps-1(RNAi)* treated embryos. Orange box encloses the unsignaled MS nucleus, Blue box encloses the signaled E nucleus. C) Quantification of GFP::SYS-1 nuclear enrichment in indicated dynein RNAi knockdowns. C’) Mean difference in nuclear GFP::SYS-1 enrichment from quantification in C. Error bars are SEM. D) Embryos ectopically expressing pEND-1::GFP::H2B in cells not descended from the E lineage in the first (orange) second (blue) and third (green) lineage after the founding of the lineage (the 2, 4, and 8 cell E descendants, respectively). D’) Representative embryos exhibiting WT (all lacZ*(RNAi)*-treated, left) and ectopic (*ecps-1(RNAi)*- and 2x*dhc-1(RNAi)*-treated for 1, 2, and 3 generations after E founding, respectively, right) P_end-1_::GFP::H2B. **\* - p<0.05, ** - p<0.01, *** - p<0.001**

Having established an increase in nuclear GFP::SYS-1 in dynein subunit depletion, even in cells not signaled to stabilize SYS-1, we turned to the effect of SYS-1 expression on target gene activation. Specifically, we used the endogenous promoter of WβA target gene *end-1* to drive GFP-tagged Histone 2B in order to evaluate the activity of SYS-1 immediate downstream targets as the signaled E cell established the endoderm tissue expansion from 2 to 4 to 8 cells (E+1, +2, or +3 divisions, respectively). Normally, this strain exhibits additional expression of the *end-1* promoter in <10% of embryos, a frequency which decreases further when SYS-1 is depleted via RNAi (Fig. 5D). In embryos depleted of either dynein subunits or RSA-2 by RNAi, however, we observed a different pattern. In these embryos, misregulation via dynein depletion appeared to increasingly overwhelm SYS-1 negative regulation in successive divisions, such that each generation after SYS-1 dependent specification increasingly expresses the *end-1* promoter driven endodermal reporter outside of the EMS lineage (Fig. 5D). For instance, in *dhc-1(RNAi)* we observe only 3% of embryos mis-expressing one cell division after the specification of the E cell type (E+1), which increases to 25% by the next division (E+2), and over 83% three divisions after specification (E+3).

We also investigated larval ACDs to determine the effect that extra transcriptionally active SYS-1 has on organismal development. We examined the seam cells (SCs) and distal tip cells (DTCs) because of their sensitivity to SYS-1 dependent gene activation (Baldwin, Phillips et al 2016; Baldwin and Phillips 2018; Lam and Phillips 2017; Mila, Putzke et al 2015). In wild-type animals, DTC lineages undergo asymmetric cell division during L1 to give rise to two DTCs, while *sys-1* mutants and animals overexpressing SYS-1 show loss or gain of DTCs, respectively, due to symmetric cell division (Fig 6A) (Miskowski, Kimble et al 2001; Siegfried, Kimble et al 2004; Kidd, Kimble et al 2005; Chesney, Kimble et al 2009). To further sensitize these cells for changes in SYS-1 dependent cell fate changes, we examined strains containing additional, transgenic SYS-1 driven by the heat shock promoter. In animals that are not subjected to heat shock, dynein depletion only rarely induced development of more than the endogenous 2 distal tip or 16 seam cells (Fig. 6B, 7B). In heat shocked animals, excess SYS-1 protein can overwhelm canonical SYS-1 regulation to develop as many as 6 DTCs (Fig. 6D). While heat shock-induced overexpression is sufficient to cause a notable shift towards the signaled cell fate, dynein subunit depletion was able to significantly enhance the effect of heat shock (Fig. 6C). Depletion of DHC-1 or ECPS-1 at the time of the somatic gonadal precursor division was sufficient to induce rare but wild-type-unobserved additional DTCs (0.3% of animals for these 2 treatments, N= 394, 363, respectively). Adding heat shock-overexpressed SYS-1 enhanced the penetrance of ectopic DTCs, with DHC-1 and ECPS-1 depletion resulting in populations where 11.9% and 6.8% of animals, respectively, develop more than 2 DTCs. Depletion of RSA-2, uncoupling SYS-1 from centrosomes, and depletion of the dynein light chain DYLT-1 further enhanced these misspecifications, increasing the proportion of the population with ectopic DTCs from 26.6% in wild type to 33.6% or 51% after RNAi depletion, respectively (Fig. 6D).

**Figure 6:**
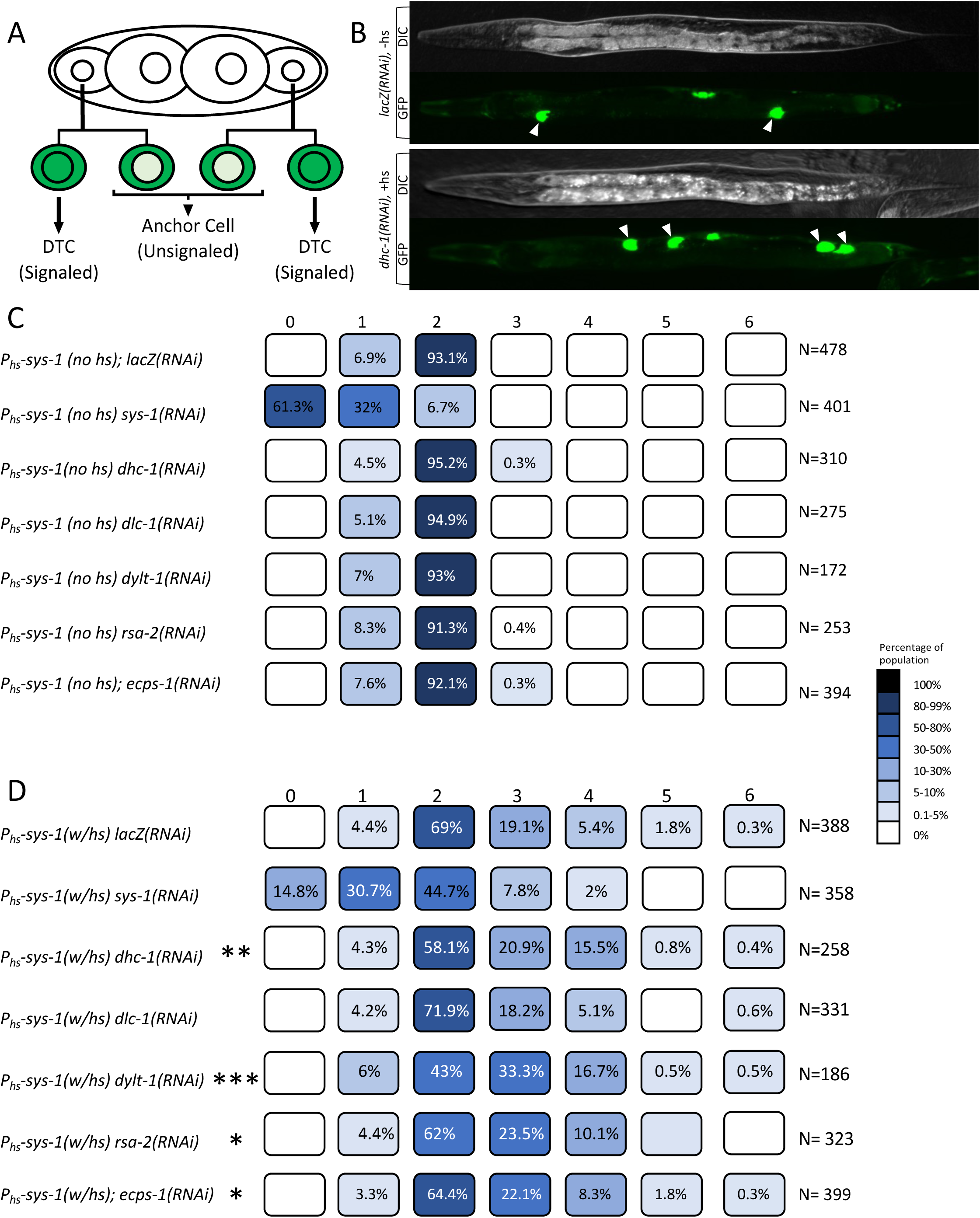
SYS-1 dysregulation via dynein knockdown can cause rare distal tip cell fate conversions. A) Diagram of *C. elegans* distal tip cell signaled lineage specification in the somatic gonadal precursor. B) Representative image of WT (top) and ectopically induced (bottom) distal tip cells (DTCs). To distinguish DTC’s from pLAG-2 driven vulval expression, DTCs are marked with white arrowheads. C) Number and relative frequency of DTCs via anatomical marker in untreated animals and in backgrounds with RNAi depletion of various dynein subunits. D) Number and relative frequency of DTCs marked by anatomical marker when SYS-1 is overexpressed via a heat shock-driven transgene in both untreated animals and in backgrounds with RNAi depletion of various dynein subunits. * = p<0.05, ** = p<0.01, *** = p<0.001 by one-sided Mann-Whitney test

**Figure 7:**
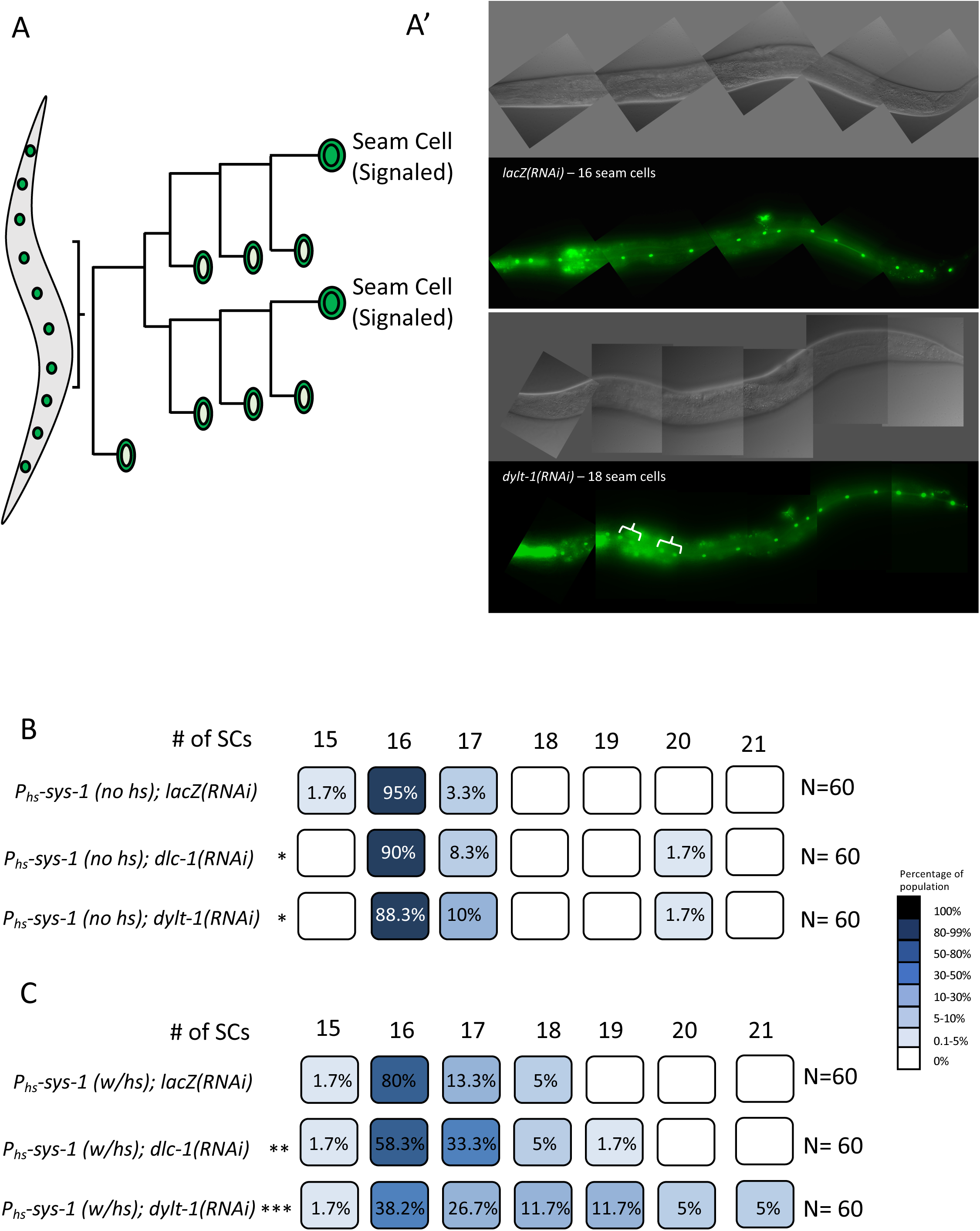
SYS-1 dysregulation via dynein knockdown can cause rare seam cell fate conversions. A) Diagram of *C. elegans* seam cell (SC) signaled lineage. B) Representative image of WT (top) and ectopically induced (bottom) SCs. Likely ectopic SC exhibit spatially distinct ‘doublets’, highlighted here in brackets. C) Number and relative frequency of SCs via anatomical marker when in untreated adults and in backgrounds with RNAi depletion of various dynein subunits. D) Number and relative frequency of SCs via anatomical marker when SYS-1 is overexpressed via a heat shock-driven transgene in both untreated larvae and in backgrounds with RNAi depletion of various dynein subunits. * = p<0.05, ** = p<0.01, *** = p<0.001 by Mann-Whitney test

The SC lineage also undergoes several WβA-dependent ACDs throughout larval development (Fig 7A, B). Similarly, in the SC tissue, depletion of dynein light chains alone is sufficient to increase the frequency of ectopic SCs from 3.3% of larvae with ectopic SCs in control RNAi conditions to 11.7% or 10% of animals as seen in DYLT-1 and DLC-1 depletions, respectively (Fig. 7C). These misspecifications are further exacerbated by SYS-1 overexpression, wherein more than 60% of the *dylt-1(RNAi)* larvae have greater than wild type seam cell numbers, developing as many as 21 SCs, (Fig. 7D). These observations collectively support SYS-1 centrosomal enrichment by dynein as a mechanism to reinforce SYS-1 function in specifying SYS-1-dependent cell fate decisions. Therefore, compromising the centrosomal SYS-1 regulatory system by dynein depletion is rarely sufficient to induce misspecifications in otherwise wildtype animals, but notably enhances the effects of ectopic SYS-1 accumulation. Therefore, dynein-dependent centrosomal regulation of transcriptional coactivator SYS-1 increases the robustness of WβA-regulated cell fate decisions after ACD.

## Discussion

Precise control of cell fate determinants is critical for normal development and homeostasis, particularly in the rapidly dividing cells of the early embryo. Regulation of these factors includes the distribution and inheritance of these determinants by daughter cells. Here we demonstrate that centrosomal negative regulation of *C. elegans* β-catenin SYS-1 is enhanced during asymmetric division by trafficking to centrosomes by the microtubule motor dynein. Depletion of individual dynein subunits or associated proteins is sufficient to reduce the enrichment of centrosomal SYS-1, and temporally specific ablation of dynein function by *dhc-1(ts)* was sufficient to remove observable SYS-1 nearly completely and disproportionately from centrosomes. However, some methods of disrupting dynein/microtubule trafficking result in an unexpected increase in centrosomal accumulation of SYS-1 protein. These increases, caused by microtubule disruption via nocodazole and depletion of SYS-1 negative regulator adapter ECPS-1, phenocopy the increased centrosomal enrichment and immobile fraction of SYS-1 seen in *rpt-4(RNAi)* depletion. We therefore propose a model wherein both SYS-1 and regulatory components of the ubiquitin-proteasome system, are dependent on microtubule-mediated trafficking to centrosomes by the dynein complex in order to appropriately reinforce SYS-1 asymmetric regulation across daughter cells (Fig. 8).

**Figure 8:**
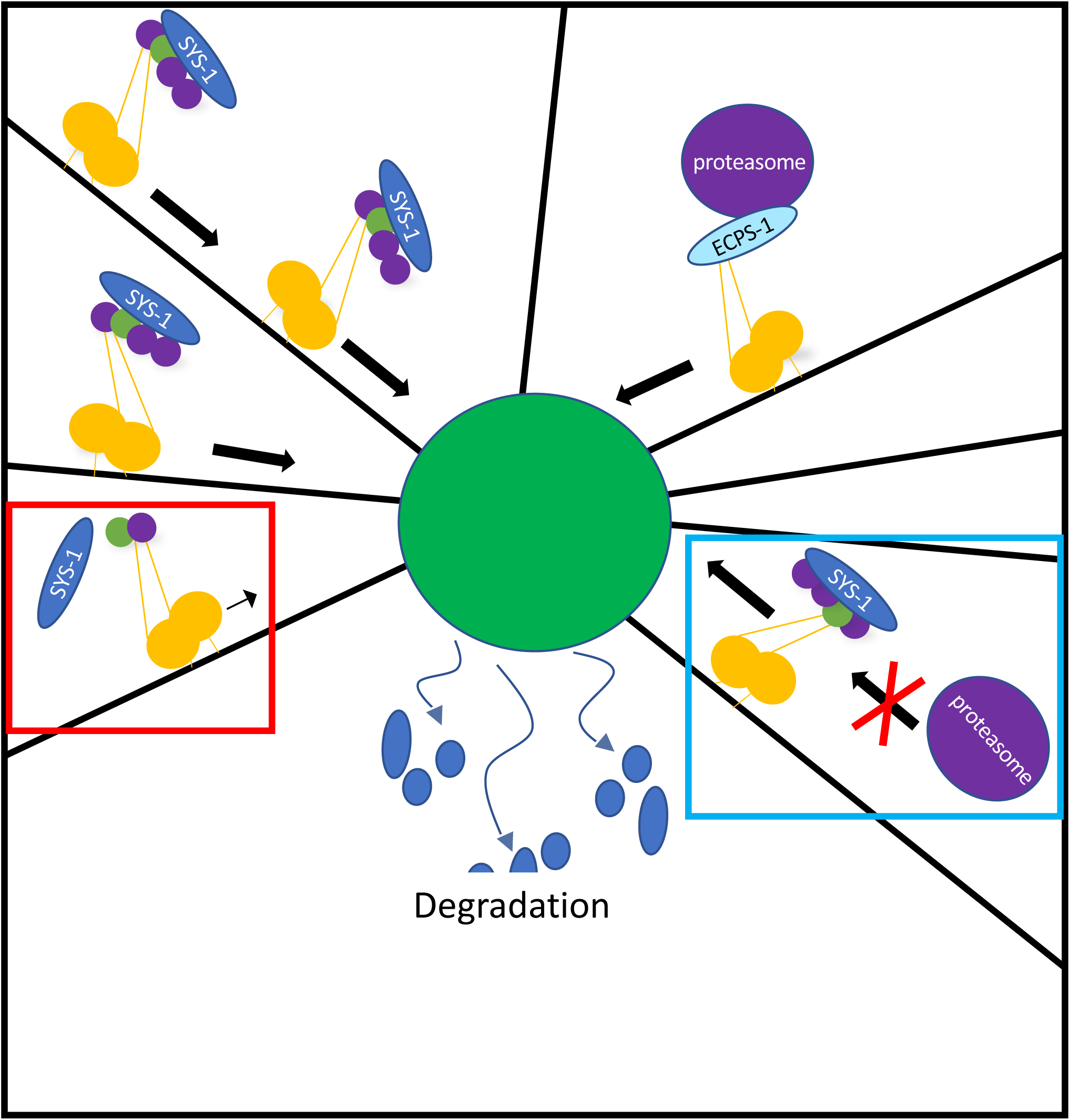
Proposed model. Speculation into the various processes by which SYS-1 and its regulators appear to be localized to mitotic centrosomes. Both SYS-1 and some component of its centrosomal proteasome-dependent negative regulators, here represented by purple circle, demonstrate some dependence on microtubule mediated trafficking. While various dynein subunits are requisite components of a functional dynein complex, the negative effect of their depletion on SYS-1 centrosomal steady state indicates a preferential or more frequent trafficking of SYS-1. Relatively rare proteasome trafficking events mediated by ECPS-1 meanwhile, are sufficient to maintain appropriate SYS-1 centrosomal enrichment. This less efficient proteasomal trafficking, however, renders it vulnerable to nocodazole-mediated decrease in overall microtubule trafficking/network formation efficiency. Red inset depicts decreased processivity or cargo binding capacity induced by dynein light chain RNAi depletions. Blue inset depicts depletion of adapter ECPS-1 and resulting depletion of proteasome-trafficking specific dynein complexes, while not otherwise affecting the microtubule motor.

In this model, SYS-1 enriches at centrosomes and is degraded or marked for degradation before leaving the centrosome. Rather than a static centrosomal enrichment, this model entails continual trafficking by microtubule motors to maintain an observable centrosomal pool of SYS-1. The observable CEI then measures a steady state of centrosomal enrichment at centrosomes, wherein SYS-1 is localized to centrosomes more rapidly than it is marked and cleared. SYS-1 therefore becomes visibly enriched at centrosomes compared to cytoplasmic levels. This model predicts that even subtle disruptions to microtubule mediated trafficking could limit the ability of cells to maintain wild type SYS-1 centrosomal enrichment, assuming the SYS-1 clearance rate remains unchanged.

Interestingly, the *rsa-2(RNAi)* enhancement of kinetochore localization of SYS-1 in this model is consistent with ‘traffic jams’ observed in microtubule mediated trafficking (Leduc, Howard et al 2012). In these cases, crowding motors by increasing the local concentration of motor proteins, introducing microtubule-binding obstacles, or, importantly for our model, preventing terminal motor dissociation limits the activity of motor proteins (Nam and Epureanu, 2017; Ferro, Yildiz et al 2019). Evidence of traffic jams in vivo have been shown in neuronal microtubule transport, as neuronal kinesin and its cargo synaptotagmin accumulate when trafficking is disrupted by hypomorphic kinesin alleles. The disrupted motor complexes then accumulate upstream of the trafficking impediment, in an accumulation of both motor and cargo (Hurd and Saxton 1996).

The kinetochore localization of SYS-1 is consistent with a traffic jam model, thus making SYS-1 trafficking by microtubule motors a more attractive model. Decreasing centrosomal SYS-1 localization by *rsa-2(RNAi)* increases SYS-1 protein in resulting daughter cells (Vora and Phillips 2015). A severely limited centrosomal recruitment of SYS-1 in *rsa-2(RNAi)* may also suggest problematic SYS-1/motor protein unloading upon reaching the MTOC centrosome. Similarly, P_pie-1_::GFP::SYS-1, which appears overexpressed compared to P_sys-1_::VENUS::SYS-1, may exacerbate SYS-1 traffic jams by increasing the cytoplasmic SYS-1 pool. In either case, the combination of P_pie-1_::GFP::SYS-1 and *rsa-2(RNAi)* results in a further increase in the incidence of SYS-1 localization to kinetochore microtubules, suggesting increased SYS-1 levels coupled with decreased centrosomal capturing may reveal an otherwise transient pool of SYS-1 en route to the MTOC centrosome. Inefficient SYS-1 capture in *rsa-2(RNAi)* embryos likely further crowds SYS-1-containing microtubule motor complexes into ‘traffic jams’ because, in embryos lacking a mechanism to anchor SYS-1 to the centrosome, SYS-1 is trafficked toward centrosomes more quickly than it is captured there and processed. This shift results in an observable SYS-1 accumulation at regions of high microtubule occupancy, such as kinetochore microtubules and the cell cortex, both of which are increased after RSA-2 depletion.

Conversely, centrosomal protein processing by either post-translational modifications or degradation via the proteasome likely requires the trafficking of fewer, yet reusable enzymes. The disparity between substrate vs enzyme trafficking requirements would suggest that clearance of centrosomal substrates is less vulnerable to mild or short-term disruptions to trafficking than the maintenance of a centrosomal substrate pool. Upon long-term or severe disruptions to microtubule trafficking, a trafficking requirement for clearance mechanisms becomes apparent. Our model predicts that disruption of either SYS-1 localization (inefficient trafficking to the site of regulation) or disrupting the localization of a member of the Ubiquitin-Proteasome System (UPS) (where SYS-1 is trafficked and accumulates, but centrosomal regulation/degradation is ineffective) both result in increased SYS-1 dependent gene activation and differentiation in daughter cells, consistent with excessive inheritance. While this system of centrosomal localization and clearance of SYS-1 does not appear necessary for most WβA differentiation events, disruptions to SYS-1 trafficking or processing limit the system’s ability to tolerate perturbations. That is, SYS-1 centrosomal localization generates robustness in SYS-1 dependent cell fate decisions (Fig. -7) due to parallel trafficking mechanisms; disrupting either trafficking mechanism results in excessive SYS-1 dependent activity in daughter cells of mothers with either increased or decreased SYS-1 CEI.

Why does no single dynein disruption treatment fully recapitulate the *rsa-2(RNAi)* phenotype? While we see individuals within several dynein knockdown treatments with nearly *rsa-2(RNAi)*-like SYS-1 centrosomal uncoupling, these populations also include embryos with roughly wild type centrosomal SYS-1 enrichment. However, given the severely deleterious nature of dynein subunit loss-of-functions (Sönnichsen, Neumann et al 2005) and the variability of feeding RNAi knockdowns as compared to null mutant strains (Tavernarakis, Driscoll et al 2000), it seems likely that our significant decreases in SYS-1 centrosomal recruitment represent a phenotypically incomplete knockdown. Surviving embryos then only exhibit hypomorphic dynein function, insufficient to induce catastrophic defects in spindle assembly but sufficient to affect SYS-1 centrosomal enrichment. However, it is also possible that there is partial redundancy between the SYS-1 relevant functions of the dynein motor. Introducing *dhc-1(ts)* overcomes either incomplete knockdown or redundancy since a thorough dynein disruption can robustly uncouple centrosomal SYS-1.

Why, then, do we observe an increase in centrosomal SYS-1 steady state upon either long term lower dose nocodazole-mediated knockdowns of trafficking or by RNAi depletion of trafficking subunit ECPS-1? In response to these data (Fig. 3), we propose the additional trafficking of a SYS-1 negative regulatory component to this model. Disrupting the trafficking of centrosomal SYS-1 negative regulators should result in a previously unseen increase in centrosomal SYS-1 steady state, while exhibiting a similar effect on SYS-1 retention in daughter cells, downstream gene expression, and cell fate decisions (Fig. 3-7). However, the difference in SYS-1 enrichment between most dynein depletions and that of ECPS-1 or nocodazole treatment suggests a differential efficiency in trafficking or centrosomal capture rate between SYS-1 and its negative regulators, such as the proteasome.

Most disruptions to the dynein motor that reduce SYS-1 centrosomal accumulation, mediated by maternally fed RNAi, are consistent but likely incomplete because of their requirement for oogenesis and early embryonic viability. Relatively mild dynein knockdowns resulting in viable embryos are likely able to form dynein complexes, but at a far reduced rate. These dynein-limited embryos should therefore decrease the rate of continual SYS-1 trafficking while not preventing the centrosomal localization of an occasional regulatory components, perhaps of the Ubiquitin-Proteasome System (UPS). The reusable tagging or proteolytic machinery is likely sufficient to enzymatically process SYS-1 that still localizes to the centrosome, but the rate of incoming SYS-1 is significantly reduced. Similarly, depletion of dynein function by *dhc-1(ts)* effectively depletes all dynein function but can only do so for brief portions of mitosis without inducing mitotic spindle collapse.

While dynein depletions limit the volume of trafficking by limiting the number of properly constructed dynein motors, they do not limit the ability of an individual, functional dynein motor to traffic. The experimental conditions that increase the steady state of centrosomal SYS-1, nocodazole and *ecps-1(RNAi)*, instead limit the scope of dynein cargo trafficking. ECPS-1, putative adapter between dynein and the proteasome, increases the steady state of both SYS-1 and additional centrosomal proteasome target ZYG-1. Increased enrichment of both proteins is consistent with the conserved role of ECM29 for ECPS-1, and results in embryos that are not inhibited for overall dynein function but limited in their ability to traffic proteasome. Nocodazole treatment similarly does not directly affect the processivity of dynein complexes that form on the comparatively limited microtubule network. However, it does severely affect the reach of the microtubule network itself (Fig. 3A). Our data therefore suggests that the proteasome requires less continual reinforcement by microtubule-mediated trafficking but does require more extensive microtubule formation than is allowed in our nocodazole-treated embryos.

A role for trafficking in both SYS-1 and its regulatory components, then, suggests a differentially maintained steady state interaction for the proteasome and centrosomal SYS-1. While SYS-1 is rapidly and continuously trafficked, it is also rapidly processed by centrosomal regulatory proteins. That is, the apparently rapid processing of centrosomal SYS-1 also requires rapid dynein-mediated flux to maintain its centrosomal steady state. Any disruption to dynein processivity will therefore allow slower SYS-1 centrosomal enrichment to be overcome by its centrosomal processing, reducing the SYS-1 CEI by slowing the transfer of SYS-1 from cytoplasm to centrosome for degradation. By comparison, the proteasome or ubiquitin ligases endure at the centrosome, continually processing centrosomal targets. Our proposed model therefore predicts that short term or partial disruptions to the dynein motor (Fig. 8, red inset) allow for the eventual establishment of a centrosomal proteolysis pathway, which may affect both SYS-1 and proteasome localization, but nevertheless result in decreasing steady state levels of centrosomal SYS-1 (and presumably other proteasome targets that require active trafficking) as occasional proteasome centrosomal localization overcomes the impaired localization of SYS-1. Conversely, long term systemic disruptions to microtubule-mediated motors like that seen in nocodazole treatment may still capture SYS-1 in their comparatively limited microtubule network, while being disproportionately deprived of the larger proteasome complex (Fig. 8, blue inset).

Our model that both SYS-1 and its regulators are trafficked was supported but the identification of a trafficking component that affects one but not both arms of our trafficking model. Most of our perturbations, generally affecting components of the dynein machinery, may exert their effects by either detaching SYS-1 from the motor or by sabotaging all function of the motor complex. However, the specificity of our two transport mechanisms was exhibited by one dynein-protein interaction, via depletion of ECPS-1/the putative ECM29 subunit of the motor. Unlike the other subunits presenting strong SYS-1 phenotypes, ECM29 has not yet been implicated in dynein function beyond its adaptor role but has notable effects on proteasome localization and response to stress (Gorbea, Rechsteiner et al 2010; Haratake, Chiba et al 2016). Because *ecps-1(RNAi)* phenocopies the centrosomal GFP::SYS-1 accumulation seen in proteasome knockdown via *rpt-4(RNAi)* as well as microtubule network disruption via nocodazole treatment, we propose that these observations support a role for proteasomal localization as well as function for proper centrosomal clearance of SYS-1 (Fig. 8).

While this model and our experiments to test it have been largely centered on SYS-1/β-catenin, many developmental and temporally regulated proteins are known to pass through the centrosome (Vora and Phillips 2016; Brown, Welch et al 1994; Johnston, Kopito et al 1998; Huang and Raff 1999; Reck-Peterson, Carter et al 2018), as the centrosomal proteasome is far from SYS-1-specific. It is not unreasonable, then, to presume that other proteins may benefit from dynein trafficking to something as short-lived and yet influential to daughter cell inheritance as the mitotic centrosome. Particularly early in eukaryotic development, subtle changes in the inheritance of cell fate determinants like SYS-1 can have a profound effect on the ability of an organism to respond to environmental insults. For example, the asymmetric nuclear localization of SYS-1 indicative of differential cell fate has been shown to arise early, less than 70 seconds after the establishing division (Baldwin and Phillips 2014; Vora and Phillips 2015). We therefore propose that an active dynein mediated trafficking, at least for cell fate determinants like SYS-1, could provide a universal buffering against excessive inheritance and consequentially rare but deleterious cell fate changes.

## Materials and Methods

### Strains

Strains were maintained on Op50-inoculated Nematode Growth Media (NGM) plates at 15° (BTP51, BTP220) or 25° C (TX964, BTP51, LP563, TX691) using typical *C. elegans* methods. These strains contained the following transgenes and alleles: **N2** (wild type); **TX964** (unc-119(ed3) III; him-3(e1147) IV; teis98 [P_pie-1_::GFP::SYS-1]); **BTP51** (unc-119(ed3) III; tjIs71 [pie-1 promoter::mCherry::H2B, pie-1 promoter::2x mCherry::tbg-1, unc-119(+)]; qIs95 III) ; **BTP220** (unc-119(ed3) III; tjIs71 [pie-1 promoter::mCherry::H2B, pie-1 promoter::2x mCherry::tbg-1, unc-119(+)]; qIs95 III; dhc-1(or195) I.) ; **TX691** (unc-119(ed3) III; teIs46 [pRL1417; end-1p::GFP::H2B + unc-119(+)]) ; **OC341 (**unc-119(ed3) III; bsIs8[pMS5.1:unc -119(+) pie -1- gfp-zyg-1C-terminus]); **BTP93** (*wIs78[SCMp::GFP + ajm-1p::GFP + F58E10* (cosmid) *+ unc-119(+*)] *IV; uiwEx22*[*pHS::sys-1*]); **BTP97** (*qIs56[lag-2p::GFP + unc-119(+)] V;uiwIs3[pHS::sys-1*]); **AZ235** (*unc-119(ed3) III; ruIs48[P_pie-1_::gfp::tbb-1*]); **TH107 (**P_pie-1_::RSA-2:: GFP)

### RNAi

To perform RNAi knockdown of target genes, we used HT115 bacteria containing the pL4440 plasmid with a T7-flanked target gene insert. Most of these were obtained from a library supplied by the Ahringer lab via Addgene (Kamath and Ahringer 2003). Those which were otherwise obtained (dlc-1) used target gene-flanking primers with *C. elegans* cDNA to generate a new insert. These bacteria were used to seed Isopropyl β-d-1-thiogalactopyranoside (IPTG)-containing, inducing plates as described previously. Worms were generally plated to RNAi after Sodium Hypochlorite Synchronization as L1’s. More deleterious knockdowns (*dlc-1, dhc-1)* were washed from OP50 plates at 24 hours before imaging (L3/L4) or 12 hours before imaging (late L4/early adult) (Timmons 2006; Bekas and Phillips 2020). DLC-1 was derived using RT primers and digested with Xho I and HIND III.

### Compound Microscopy, CEI, and Image Processing

To obtain samples, mothers were anesthetized with 250 μM levamisole and dissected *via* a clean surgical blade. Embryos were then transferred to a 2% agarose pad and imaged on a Zeiss Axio Imager.D2. All exposure times were kept constant. Centrosomal measurements were collected from the mitotic P1 cell of the early embryo during anaphase, at approximately similar inter-polar distance (Fig. S5). Images were exported as uncompressed TIF’s, and fluorescence intensity was determined by measuring the mean fluorescence of centrosomal puncta in ImageJ. The mean intensity of the remaining embryonic cytoplasm was subtracted from the centrosomal mean to produce the CEI measurement, and identical measurements on 2x mCherry::tbg-1 were used for TEI. END-1 family nuclei were assayed via the TX691 strain. GFP-tagged Histone intensity was measured relative to the cytoplasmic autofluorescence.

### Temperature Sensitive Allele Imaging

BTP51 and BTP220 were maintained together at 15°C and were imaged as above. Ambient temperature in the microscope room was decreased to ∼18°C for ‘permissive’ imaging and upshifted to ‘restrictive’ 26°C by moving slides to a Pecon Tempcontroller 2000-2 heated stage. Embryos were imaged at the two temperatures sequentially, with approx. 45 seconds between upshift and image, or were imaged as populations at either the permissive or restrictive temperature.

### Nocodazole Treatment

In order to treat embryos before formation of the drug-impermeable eggshell, we exposed late L4’s to a low dose (approx. 83 μM, 25 μg/mL) nocodazole dissolved in 2% dimethyl sulfoxide (DMSO) for 16 hours. Embryos were then concentrated by one minute centrifugation at 1000xG and imaged as described above. Permeabilization of embryos was performed as described in Carvalho, Oegema et al 2011. Imaging was done as described above, but included 4% sucrose and 0.1M NaCl to the 2% agar pads to bolster embryo stability (Greenan, Hyman et al 2010; Gönczy, Schnabel et al 1999).

### Confocal Microscopy and Fluorescence Recovery After Photobleach

Confocal microscopy was performed via a Leica SP8 HyD detector system. The objective used was a 63x HC PL APO CS2 objective with N.A. = 1.4, utilizing type F immersion oil. Each image analyzed consisted of the sum of 35 z-images across the embryo, each slice approximately 0.35 µm thick. FRAP was performed on the same system. Each embryo was imaged twice before bleaching, beginning as soon as possible after Nuclear Envelope BreakDown (NEBD). The ROI, an approximately 5μm square, was then bleached at an intermediate focal plane with 100% laser power for 100 iterations. Images were taken as above, once every 7 seconds. Fluorescence intensity was measured on summed slice Z-projections in FIJI via the mean fluorescence intensity of ROIs drawn around relevant locales – either centrosome or the entire dividing cell. Recovery was assayed by subtracting the ROI post-bleach intensity from each measurement and evaluating it as a percentage of the pre-bleached centrosomal intensity.

### Heat Shock-Induced SYS-1 Overexpression

BTP93 and BTP97 worms were exposed to brief 33℃ heat shocks at relevant developmental timepoints to drive SYS-1 overexpression concurrently with signaled cell fate specification. For DTCs, BTP97 was subjected to 1-hour heat shock, 30-minutes recovery, and 30-minutes additional heat shock beginning 11.5 hours after plating. For SCs, BTP93 was subjected to 2-hour heat shock, 30-minute recovery, and additional 1-hour heat shock beginning 26 hours after plating. Both tissue types were assayed 50 hours after plating.

## Acknowledgements

We thank Kimberly Bekas and Amy Clemons for helpful comments on the manuscript. Strains were provided by the Caenorhabditis Genetics Center, which is funded by the National Institutes of Health (NIH) Office of Research Infrastructure Programs [P40 OD01440], or gifts from the laboratory of Tony Hyman (TH107) and Kevin F. O’Connell at the NIDDK (OC341). This work was supported by NIH award NIH GM114007 (B.T.P.).

## Notes

### Competing Interest Statement

The authors have declared no competing interest.

### Summary of Updates

Figure 5 panel names edited; text edited for clarity; references edited

