## Supplemental data for "Centrosomal Enrichment and Proteasomal Degradation of SYS-1/β-catenin Requires the Microtubule Motor Dynein"

Figure S1

A

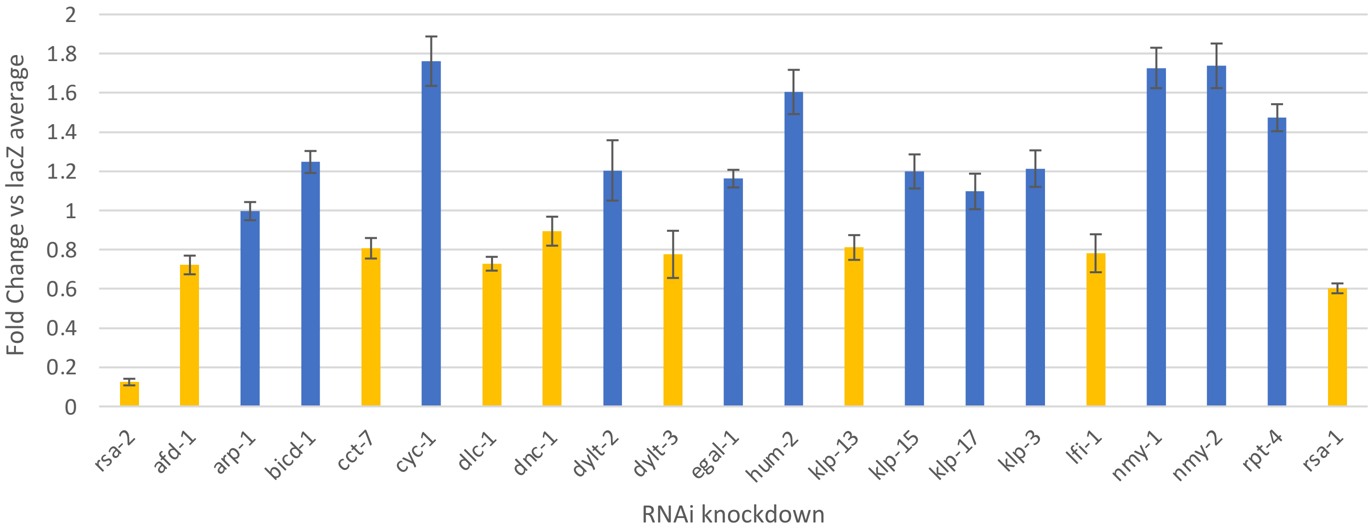


B

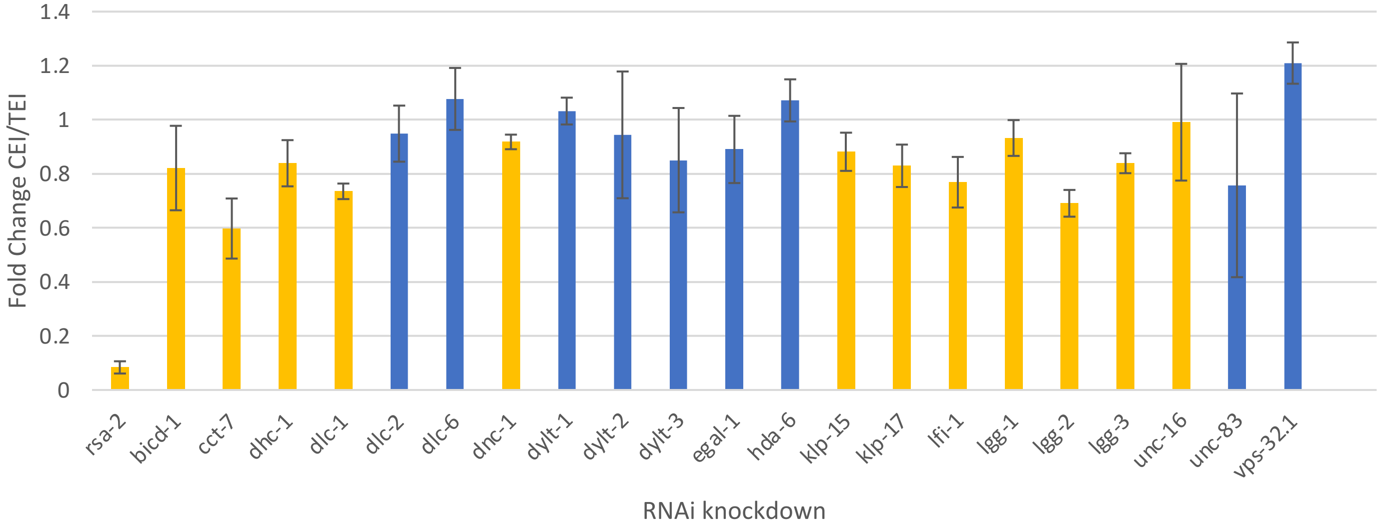


C

| Protein Function | Proteins Screened |
|---|---|
| Dynein Subunit | CHE-3, **DHC-1**, DHC-3, **DLC-1**, DLC-2, DLC-6, DLI-1, DYCI-1, DYLT-1, DYLT-2, **DYLT-3**, DYRB-1, |
| Dynein Associates | **AFD-1**, **BICD-1**, **DNC-1**, EGAL-1, LIS-1, **LFI-1**, RBA-1, RAB-7**, UNC-16**, ZYG-12, DNC-2, DNC-3 |
| Alternative Motors | BMK-1, **KLP-13**, KLP-3, KLP-7, NMY-1, NMY-2, UNC-116, HUM-2, **KLP-15**, KLP-16, **KLP-17**, KLP-20, OSM-3, UNC-104, TRAK-1 |
| Cytoskeleton Associated | ARP-11, BEC-1, GIP-1, GRDN-1, **RSA-1**, TBA-1, TBA-2 |
| Autophagy related | BEC-1, **LGG-1**, **LGG-2**, **LGG-3**, M117.3, RAB-11.1, RAB-6.1, RAB-7, VPS-32.1, VPS-35 |
| Misc. | CCT-1, **CCT-7**, CYC-1, DAF-19, EPI-1, EVL-20, FBXA-171, FBXB-54, GIPC-1, GIPC-2, HDA-6, HTT, KCA-1, MIB-1, NHR-25, SAX-2, |

**Figure S1:**

**Results of SYS-1 trafficking screen**. CEI (A) and CEI/TEI (B) depict the results, across two strains, of animals with sufficient surviving embryos to quantify. Both measurements are displayed as a fold change, relative to the mean lacZ*(RNAi)* CEI or CEI/TEI, such that a 1 value indicates an identical average CEI or CEI/TEI to negative control. Error bars are SEM. We considered partial hits, indicated in orange, treatments reducing the full range of SEM below 1. C) Simplified table of screen target functions.

Figure S2

A

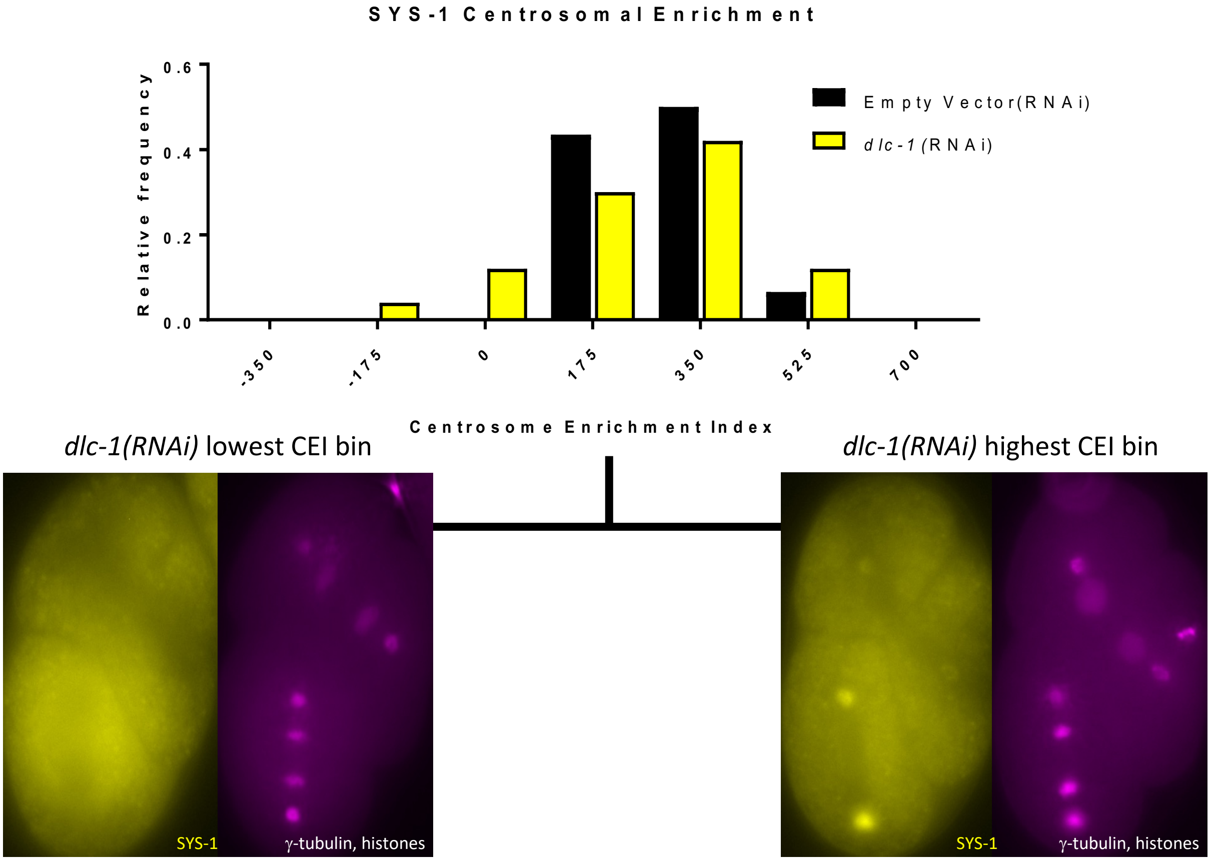
SYS-1 Centrosomal Enrichment
Relative frequency
0.0
0.2
0.4
0.6
Empty Vector(RNAi)
*dlc-1* (RNAi)
-350
-175
0
175
350
525
700
Centrosome Enrichment Index
*dlc-1(RNAi)* lowest CEI bin
*dlc-1(RNAi)* highest CEI bin
SYS-1
γ-tubulin, histones
SYS-1
γ-tubulin, histones

B

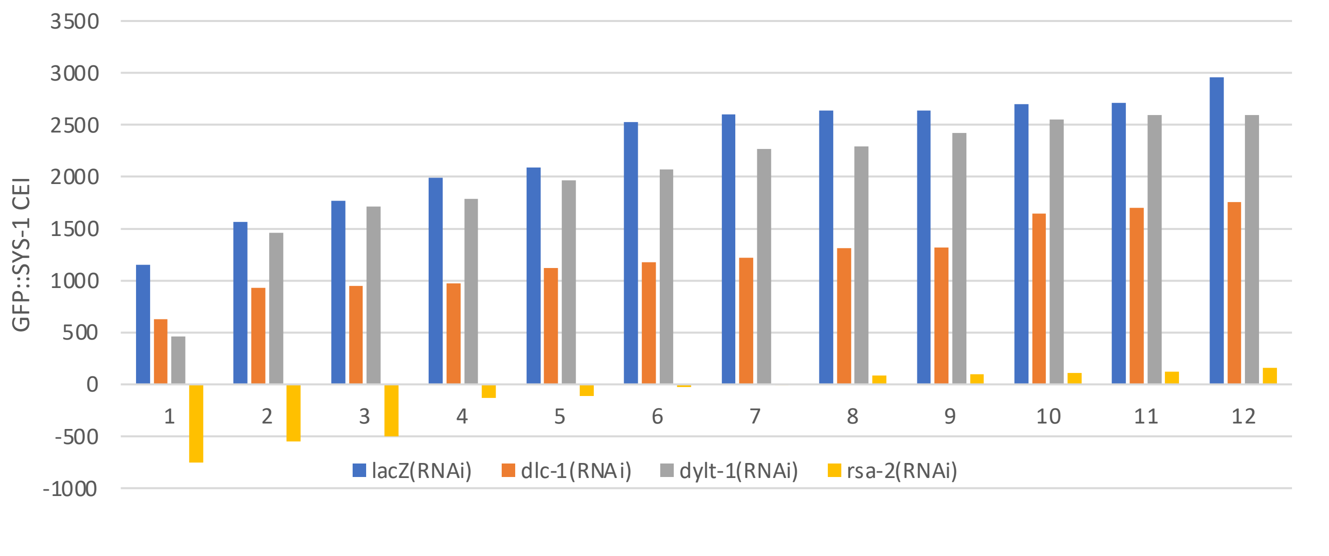
GFP::SYS-1 CEI
3500
3000
2500
2000
1500
1000
500
0
-500
-1000
1
2
3
4
5
6
7
8
9
10
11
12
lacZ(RNAi)
dlc-1(RNAi)
dylt-1(RNAi)
rsa-2(RNAi)

**Figure S2:**

**Dynein depletions by RNAi result in rare but severe decrease in SYS-1 CEI.** In individual experiments, depletion of DLC-1 by RNAi induces occasional near complete losses of fluorescently tagged SYS-1 centrosomal enrichment. A) Histogram depicting frequency of binned CEI measurements in L4440 empty vector (black) and *dlc-1(RNAi)*(yellow) populations (N=46 and 50, respectively). Images show the extent of *dlc-1(RNAi)* embryos SYS-1 CEI range, corresponding to the lowest (left) and highest (right) bins of SYS-1 centrosomal enrichment in this population. B) Twelve rank ordered embryos of the depicted dynein depletion or control exhibit a largely overlapping, but occasionally severe, range for dynein depletion effect on SYS-1 CEI.

Figure S3

A

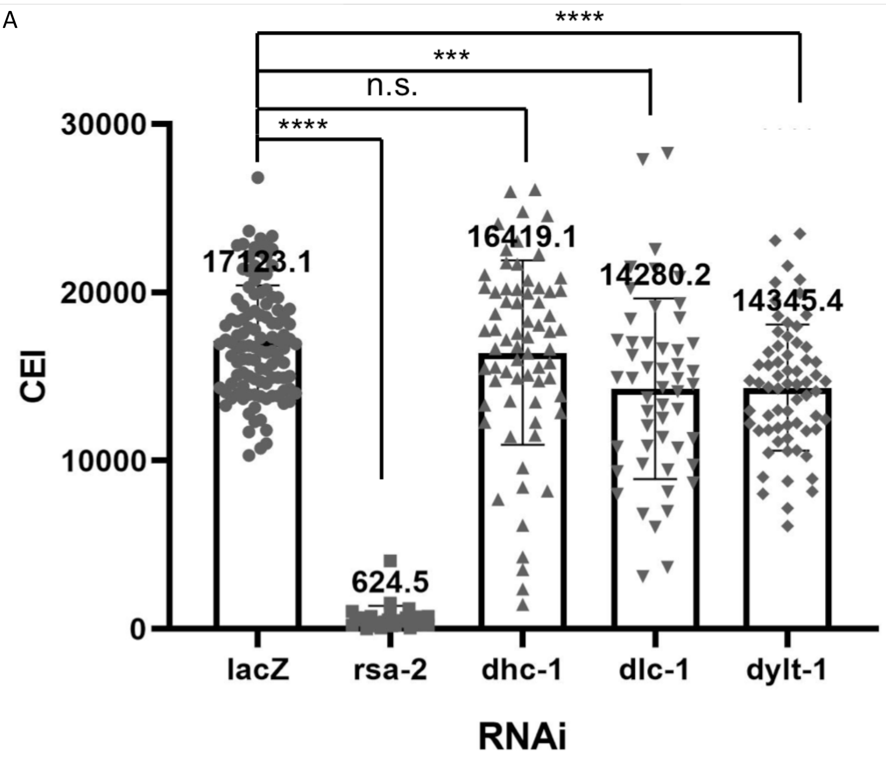

****
***
n.s.
****
30000
20000
10000
0
CEI
17123.1
624.5
16419.1
14280.2
14345.4
lacZ
rsa-2
dhc-1
dlc-1
dylt-1
RNAi

**Figure S3:**

**Dynein depletions by RNAi Slightly Reduce RSA-2 Enrichment.** Centrosomal enrichment of Ppie-1::RSA-2::GFP, measured by CEI. Each symbol represents an individual centrosome. N= 83, 29, 68, 46, 55, respectively

### Figure S4

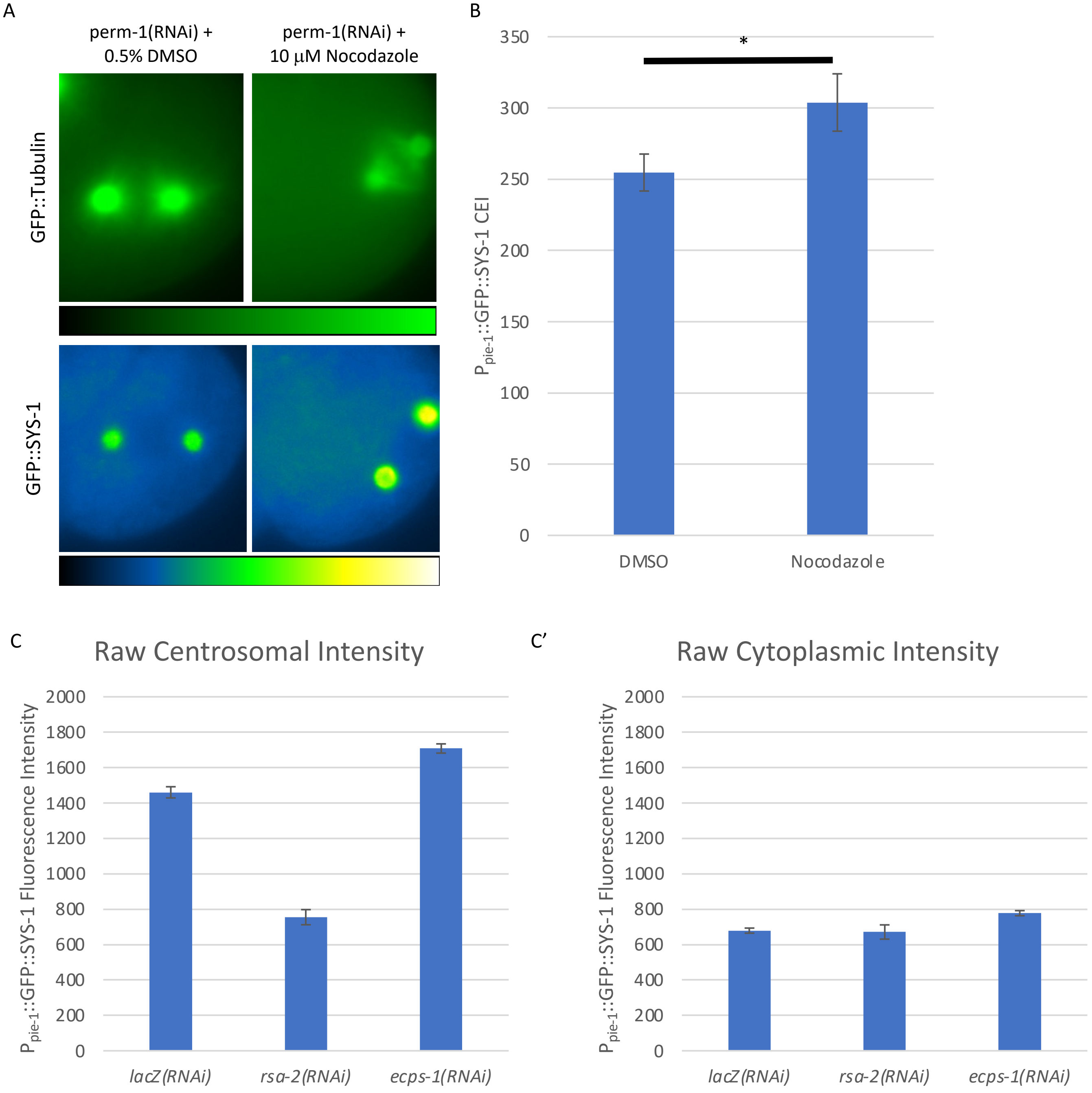
A
perm-1(RNAi) +
0.5% DMSO
perm-1(RNAi) +
10 μM Nocodazole
GFP::Tubulin
GFP::SYS-1
B
P pie-1::GFP::SYS-1 CEI
*
350
300
250
200
150
100
50
0
DMSO
Nocodazole
C
Raw Centrosomal Intensity
P pie-1::GFP::SYS-1 Fluorescence Intensity
2000
1800
1600
1400
1200
1000
800
600
400
200
0
lacZ(RNAi)
rsa-2(RNAi)
ecps-1(RNAi)
C'
Raw Cytoplasmic Intensity
P pie-1::GFP::SYS-1 Fluorescence Intensity
2000
1800
1600
1400
1200
1000
800
600
400
200
0
lacZ(RNAi)
rsa-2(RNAi)
ecps-1(RNAi)

**Figure S4:**

P**ermeabilization of embryos supports Nocodazole effect on centrosomal SYS-1** A) Representative images of Ppie-1::GFP::Tubulin (top) and Ppie-1::GFP::SYS-1 (bottom) after eggshell permeabilization by *perm-1(RNAi)* Nocodazole resulted in spindle centering or rotation defects in 61% of imaged GFP::Tubulin embryos, N=11/18, suggesting defects in astral microtubules. Intensity LUT scale is provided below each set of images. CEI measure of *perm-1(*RNAi) treated Ppie-1::GFP::SYS-1 embryos in either 0.5% DMSO (DMSO) or 10μM Nocodazole (Nocodazole) N = 43, 39 respectively. B) Separated Centrosome (B) and Cytoplasmic (B') fluorescence intensity measurements of negative control *lacZ(RNAi), rsa-2(RNAi), and ecps-1(RNAi)*.

Figure S5

A

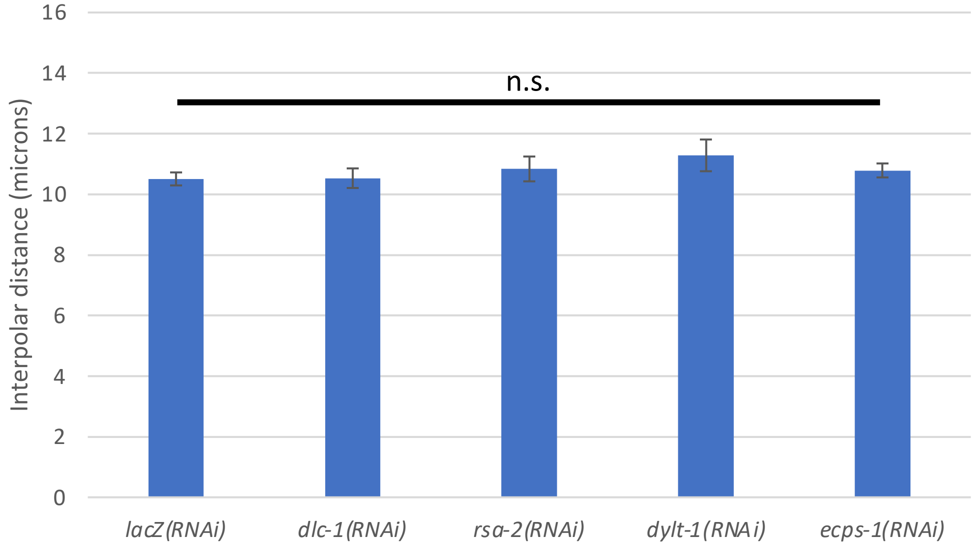

n.s.
Interpolar distance (microns)
0
2
4
6
8
10
12
14
16
lacZ(RNAi)
dlc-1(RNAi)
rsa-2(RNAi)
dylt-1(RNAi)
ecps-1(RNAi)


B

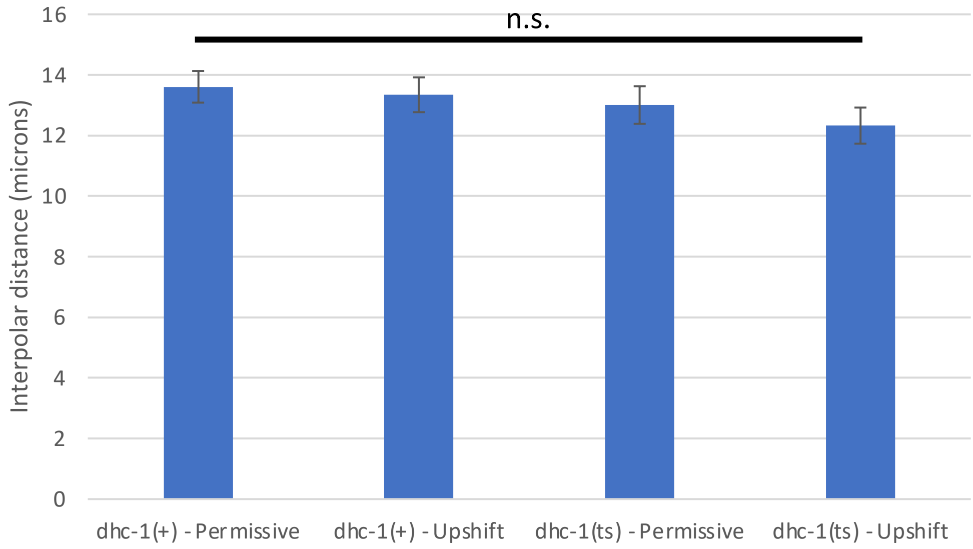

n.s.
Interpolar distance (microns)
0
2
4
6
8
10
12
14
16
dhc-1(+) - Permissive
dhc-1(+) - Upshift
dhc-1(ts) - Permissive
dhc-1(ts) - Upshift

**Figure S5:**

Dynein Knockdown Embryos Progressed Similarly Through Mitosis. Distance in microns between mitotic centrosomes in embryonic measured centrosome populations in A) dynein knockdown populations and B) Wildtype and *dhc-1(ts)* embryos at the permissive and restrictive temperatures.
